# DNA Conformational Flexibility Descriptors Improve Transcription Factor Binding Prediction Across the Protein Families

**DOI:** 10.1101/2025.07.20.665724

**Authors:** Upalabdha Dey, Venkata Rajesh Yella, Aditya Kumar

## Abstract

Precise binding of transcription factors (TFs) to specific DNA sequences is fundamental to gene regulation, yet the molecular principles underpinning TF–DNA specificity remain incompletely understood. While nucleotide sequence and DNA shape are known determinants of TF binding, the role of DNA flexibility encompassing axial, torsional, and stretching dynamics— remains largely unexplored, particularly across diverse TF families. Here, we systematically integrate experimentally and computationally derived DNA flexibility descriptors into predictive models of TF–DNA binding specificity. Through extensive analyses of large-scale in vitro datasets from HT-SELEX, SELEX-Seq, protein binding microarrays encompassing mam-malian and Drosophila TFs, we demonstrate that flexibility-augmented models consistently outperform sequence based models, and DNA shape augmented models to an extent. These improvements are robust across diverse experimental platforms, and scale of the datasets, underscoring the importance of DNA conformational dynamics in indirect readout. Quantitative analyses of position-specific flexibility contributions reveal distinct “flexibility hotspots” within transcription factor binding sites and their flanking regions. This is exemplified by structural insights into the homeodomain TF MSX1, where localized DNA bendability directly correlates with enhanced binding affinity and precise recognition specificity. Finally, leveraging in vivo ChIP-Seq and DNase-Seq data from ENCODE, we further validate that DNA flexibility substantially enhances the identification of functional TF binding sites across various TF families and cellular contexts. Collectively, current findings substantiate DNA flexibility as a fundamental element of the *cis*-regulatory code and significantly advancing predictive frameworks of gene regulatory networks.

## Introduction

Eukaryotic gene regulation hinges on precise protein–DNA interactions. Transcription factors (TFs), proteins that recognize specific DNA sequences, are fundamental to this process (Lambert et al. 2018). While TFs typically bind *cis*-regulatory elements to modulate gene expression, the mechanisms governing their recognition of cognate binding sites remain a central challenge in structural and molecular biology (Boer and Taipale 2024). Specifically, how TFs select functional consensus motifs from a vast pool of putative genomic sites is not yet fully understood (Slattery et al. 2014).

TF-specific DNA recognition extends beyond consensus sequences, involving chromatin context, epigenetic marks, and higher-order structural DNA features (Rohs et al. 2010). Quantifying TF binding specificities is thus crucial to understanding the complex energetic landscape of TF–DNA interactions. High-throughput technologies such as protein binding microarrays (PBM) and high-throughput SELEX (HT-SELEX) now enable *in-vitro* determination of binding affinities for large numbers of TFs simultaneously (Martin et al. 2023; Yang et al. 2017). A conventional approach for interpreting these datasets involves constructing position weight matrices (PWMs), which model nucleotide probabilities at each position of the binding site (Stormo 2013). While PWMs are valued for their simplicity and scalability, they neglect nearest-neighbor effects between adjacent nucleotides within the TF binding site (TFBS). To address this, recent models incorporate k-mer features to capture inter-dependencies among neighboring base pairs (Liu et al. 2025; Lai et al. 2019, 2019; Hombach et al. 2016).

Complementing sequence-based models, it is essential to consider the structural configurations formed by these inter-dependencies (Rohs et al. 2010; Biswas and Basu 2023). Although PWMs approximate base-specific contacts, TFs also recognize local three-dimensional DNA conformations arising from stacking interactions of the adjacent nucleotides (Rohs et al. 2010; Zeitlinger 2020; Zeitlinger et al. 2025). Numerous studies have shown that integrating DNA shape information—like groove geometry and helical parameters—improves the predictive power of TF–DNA recognition models across diverse TF families (Yang et al. 2017; Mathelier et al. 2016; Yella, Kumar, and Bansal 2014). Notably, TFs such as bHLH and homeodomain proteins rely extensively on DNA shape readout, benefiting significantly from its inclusion for accurate binding affinity predictions (Dror et al. 2014).

Importantly, DNA shape descriptors reflect static geometries (Rohs et al. 2009; Li, Chiu, and Rohs 2024). In contrast, B-form DNA in solution displays substantial conformational flexibility (Gupta, Bansal, and Sasisekharan 1980; Bhattacharyya and Bansal 1992), enabling a single sequence to explore a thermodynamic ensemble of meta-stable shapes that eventually stabilized during the TF binding (Heddi et al. 2006, 2010; Rosa et al. 2021). Prior structural studies have underscored the pervasive role of DNA sequence–encoded flexibility in protein– DNA co-crystal structures, often influenced by protein-induced DNA bending or nucleosomal positioning (Bhattacharyya and Bansal 1990).

Despite such evidence, the contribution of DNA flexibility to TF–DNA specificity—particularly its relevance across TF families remains poorly understood. In line, Yella et al. showed that conformational flexibility in flanking regions can influence binding affinity for select zinc-finger, bZIP, homeodomain transcription factors (Yella et al. 2018). However, with the availability of large-scale *in-vitro* TF–DNA specificity datasets, a systematic investigation of DNA flexibility as a general structural determinant is now feasible.

Here, we hypothesize that integrating DNA flexibility descriptors with sequence information improves the determination of TF–DNA specificity. To test this, we analyzed the most comprehensive *in-vitro* HT-SELEX dataset of mammalian TFs (Yang et al. 2017), alongside physiologically relevant and empirically determined flexibility descriptors, including DNaseI sensitivity, nucleosomal positioning preference (NPP), twist dispersion, twist-roll-x displacement (trx), and stiffness. These features capture axial, torsional, and stretching dynamics of TFBS conformational plasticity. We demonstrate that DNA flexibility is biologically informative in distinguishing between TF recognition strategies. Flexibility-augmented models consistently outperform sequence-only models and match or exceed the performance of shape-augmented models across diverse experimental platforms and organisms. Interestingly, these observations are robust across different organism, experimental design. Moreover, flexibility-driven feature engineering sheds light on bendability readout mechanisms employed by the TFs at both binding sites and flanking sites across multiple TF families. Importantly, the sequence dependent flexibility augmented models can accurately distinguish the *in-vivo* functional TFBS from background of genomic consensus motifs, suggesting their biological relevance.

## Results

### TF-family specific DNA-binding specificity can be improved by the DNA flexibility models

To investigate the influence of DNA bendability on TF sequence specificity, we reanalyzed most extensive HT-SELEX datasets to date, containing DNA-binding affinities for more than two hundreed mammalian TFs (Yang et al. 2017). In the dataset, preferred TF binding sites are represented as M-length DNA sequences (M-words) with the measured affinities. For numerical representation, we one-hot encoded these sequences into mono-nucleotide features, and supplemented with DNA flexibility descriptors derived from physiologically relevant di- and tri-nucleotide models (Sarkar et al. 2021; Dey et al. 2023; Murthy et al. 2024; Yella et al. 2018).

The tri-nucleotide flexibility models included DNaseI bendability (from chicken erythrocyte DNaseI-cutting experiments) (Brukner et al. 1995), and nucleosomal positioning preference (NPP) (Satchwell, Drew, and Travers 1986), along with three di-nucleotide models: twist-dispersion, twist-roll-x displacement (trx). While the role of tri-nucleotide bendability in TF-DNA specificity is well-documented (Dey et al. 2023; Murthy et al. 2024; Yella et al. 2018), the di-nucleotide features are comparatively less characterized for protein-DNA recognition mechanism (Vanaja et al. 2021). Nonetheless, their derivation from *in-vitro* assays or large-scale *in-silico* modeling highlights their potential importance in DNA recognition by TFs.

To evaluate the contribution of these flexibility features across 240 TFs (see Methods for details), we performed principal component analysis (PCA) using combined sequence and flexibility encodings. PCA revealed that, first two principle components (PC) obtained from the combination of DNA flexibility and sequence features explained greater amount of variance in the current dataset (14.8% for PC1, and 9.1% for PC2), compared to the PC obtained from sequence only features(PC1: 12.9% and PC2: 9%) (**Figure 1A**). Moreover, TFs from different families formed distinct clusters along the first two PCs. Importantly, clusters incorporating 1mer with flexibility features exhibited greater separation than those based on sequence alone (1mer). For instance, the first PC derived from flexibility augmented sequence features more effectively distinguished homeodomain and bZIP family TFs compared to only sequence, as shown in **Figure 1A (right panel)**.

**Figure 1.**
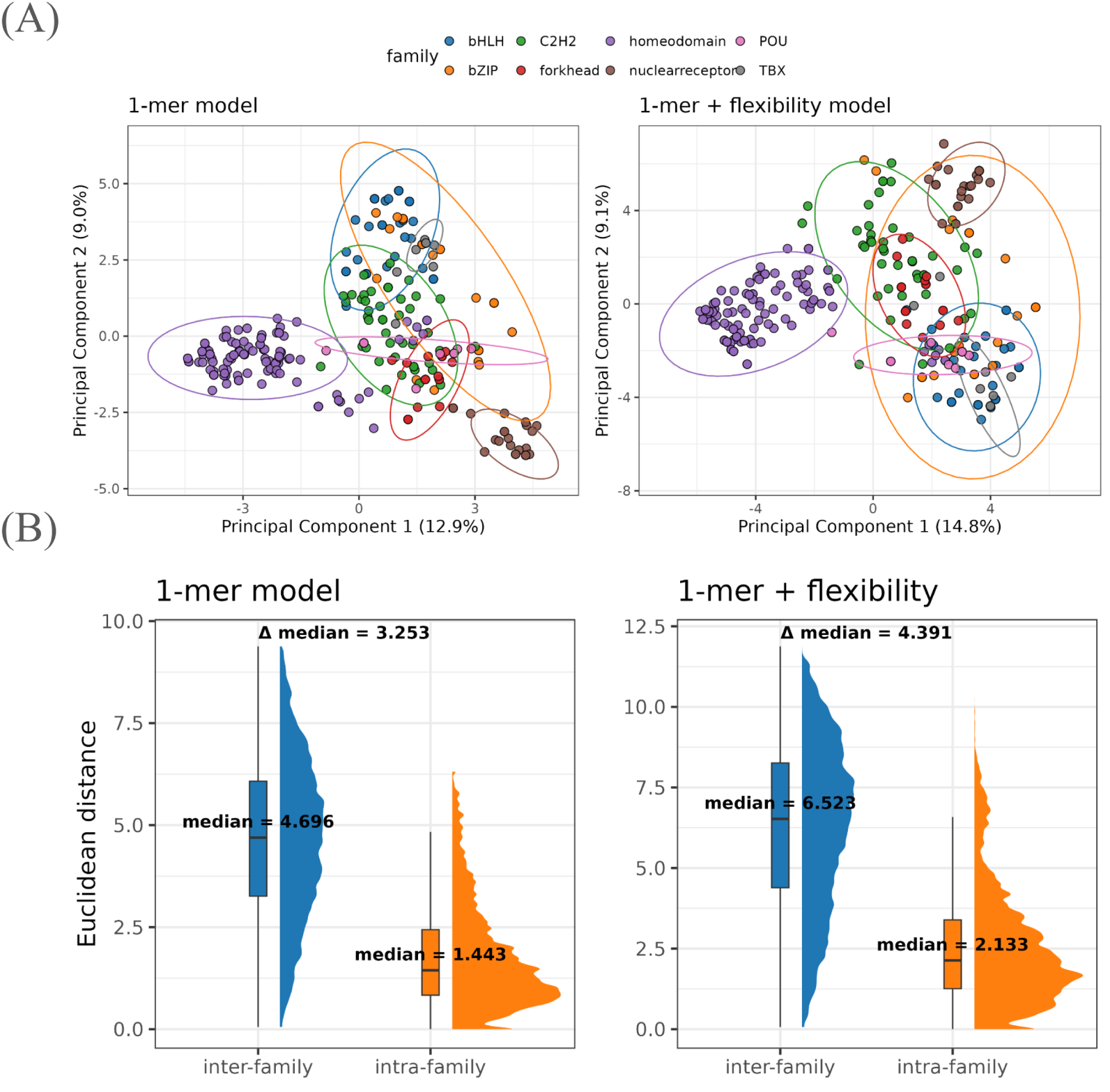
DNA Flexibility Enhances TF Family-Specific Binding Specificity. **(A)** Principal Component Analysis (PCA) of TF binding site sequences using 1-mer features only (*left*) versus 1-mer combined with DNA flexibility features (*right*). Each dot represents a TF, color-coded by its TF family. Ellipses denote the 68% confidence contour of a fitted bivariate normal distribution for each family (default in R ggplot2) **(B)** Boxplots and density distributions of pairwise Euclidean distances between TFs, calculated from the first two principal components. Plots compare models using only 1-mer features (*left*) versus those incorporating both 1-mer and flexibility features (*right*). Distances are grouped and colored by intra-family versus inter-family TF pairs.

We further quantified and compared the efficiency of flexibility features in separating the datapoints through PCA by measuring Euclidean distances between each TF pairs in the two-dimensional PC space. As shown in **Figure 1B**, inter-family distances increased markedly, while intra-family distances also increased, the median difference between the inter- and intra-family distance rise upon inclusion of DNA flexibility features.

In order to rule out the possibility that increased dimensionality alone explained the improved variance and separation, we conducted a control PCA using randomly generated flexibility features matched in mean and standard deviation to the original flexibility data (*Supplementary figure 1*). The control flexibility features with 1mer yielded lower explained variance in the datasets, and reduced Euclidean distances between the TFs from distinct families, confirming that original DNA flexibility contributes meaningful signal. Consistently, the true flexibility features increased the variance captured by the first two PCs, reinforcing their role in enhancing TF family-level discrimination.

Integrating DNA Flexibility Improves TF-DNA Specificity Prediction Beyond Sequence and Shape Models

In order to quantify contribution of DNA flexibility to transcription factor (TF) binding specificity, we adopted a quantitative modeling framework inspired by earlier studies (Zhou et al. 2015; Yang et al. 2017). Specifically, we constructed regression models to predict TF–DNA binding affinity using mono-nucleotide sequence features (1-mer) alone, and in combination with DNA flexibility descriptors (1-mer + flexibility). A consistent improvement in model performance upon incorporation of flexibility features would indicate the role of DNA flexibility in TF recognition.

We applied this approach to two in-vitro datasets: a large-scale HT-SELEX dataset profiling 215 mammalian TFs (Yang et al. 2017), and 21 SELEX-Seq datasets for eight Drosophila Hox proteins in complex with Exd cofactors (Slattery et al. 2011; Abe et al. 2015). Across 27 TF families in the mammalian datasets, flexibility-augmented models consistently outperformed the sequence-only (1-mer) models (**Figure 2A** and *Supplementary figure 2A*), highlighting the widespread role of indirect readout mechanisms.

**Figure 2.**
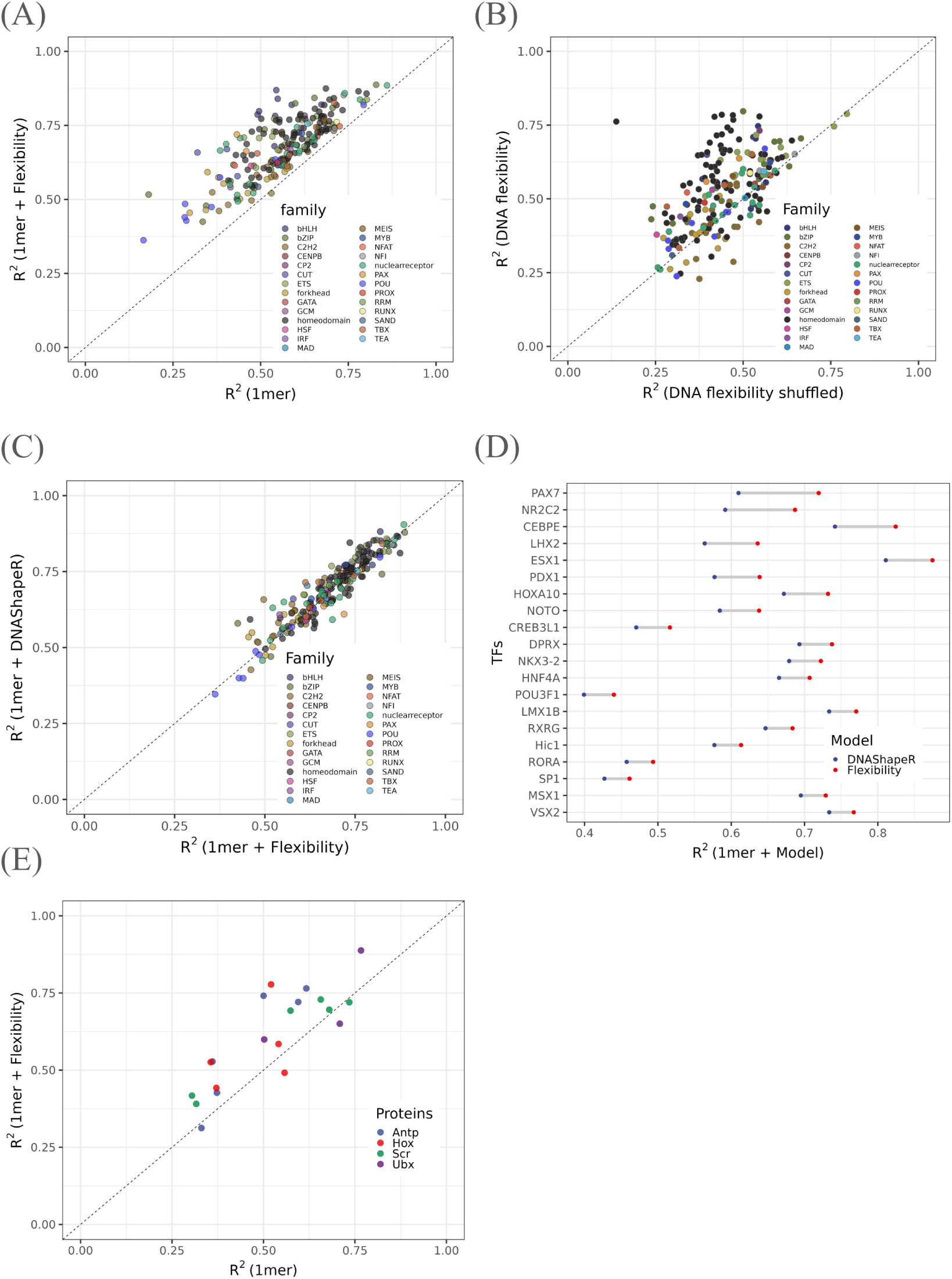
Performance Comparison of Sequence-Only and Flexibility-Augmented Models Across TF Families. **(A)** Scatter plot comparing model R² values for sequence-only (1-mer) versus flexibility-augmented models across multiple TFs colored by family. **(B)** Comparison of model performance using real versus shuffled DNA flexibility features, showing biological relevance of flexibility descriptors. **(C)** Comparison of R² values between flexibility-augmented models and DNAShape models(DNAShapeR). **(D)** Per-TF comparison of R² differences between flexibility and shape-based models highlighting cases where flexibility features outperform shape models. **(E)** Model performance comparison for Drosophila Hox proteins using sequence-only and flexibility-augmented models.

To quantify this improvement, we compared the coefficient of determination (R²) across families. Statistically significant increases in R² (Wilcoxon rank-sum test) were observed for several TF families—notably bHLH, bZIP, ETS, nuclearreceptor, and homeodomain—while C2H2 and GATA factors showed no significant gain (*Supplementary figure 2B*). It is worth noting that representation varied across TF families in the dataset, which may influence the statistical power of these comparisons. Nonetheless, over half of the TFs analyzed (107 out of 215) exhibited more than a 10% increase in predictive performance upon inclusion of flexibility features (**Supplementary Table**). As a control, we repeated the modeling using L2-regularized regression on datasets with shuffled lookup tables of the flexibility descriptors. In most cases, models trained on actual flexibility features significantly outperformed their shuffled counterparts(**Figure 2B**), confirming that improvements yielded from biologically meaningful information rather than increased dimensionality.

To further contextualize our results, we compared performance of flexibility-augmented models against DNA shape-based models (1-mer + DNAShapeR) reported by Yang et al (Yang et al. 2017). While the two approaches were generally concordant, models incorporating flexibility features achieved superior performance for several TF families, including POU, NFAT, and RUNX (**Figure 2C**). Interestingly, the bZIP factor CEBPE and the nuclear receptor NR2C2 showed the largest R² gains in flexibility-augmented models relative to shape-based ones(**Figure 2D**), reinforcing previous observations that DNaseI and NPP-derived flexibility metrics are effective predictors of TF binding (Yella et al. 2018). However, no statistically significant difference observed when comparing the models across the TF families (*Supplementary figure 2C* ).

Given the established role of DNA shape features such as minor groove width (MGW) and propeller twist (ProT) in the DNA-binding specificity of Drosophila homeodomain proteins (Slattery et al. 2011; Abe et al. 2015), we extended our analysis to SELEX-Seq data for Hox-Exd heterodimers. Among the 20 datasets analyzed, flexibility-augmented models delivered the largest performance improvements for Ubx, Scr, and Antp, mirroring gains seen with shape-based models(**Figure 2E**). While a few datasets showed better performance with sequence-only models, the overall trend supported the utility of DNA flexibility in enhancing affinity prediction for these TFs.

### DNA flexibility models robustly predict TF binding affinity across experimental platforms

Motivated by our earlier findings, we investigated whether DNA flexibility-augmented models of TF–DNA specificity could generalize beyond SELEX-based assays. To this end, we applied these models to protein binding microarray (PBM) datasets, which employ distinct experimental principles compared to exponential enrichment-based methods. Initially, we analyzed universal PBM (uPBM) data for multiple mouse TFs from the DREAM5 challenge (Weirauch et al. 2013). For the majority of TFs examined, flexibility-augmented sequence models significantly improved predictive accuracy over mono-nucleotide (1-mer) sequence-only models (**Figure 3A**).

**Figure 3.**
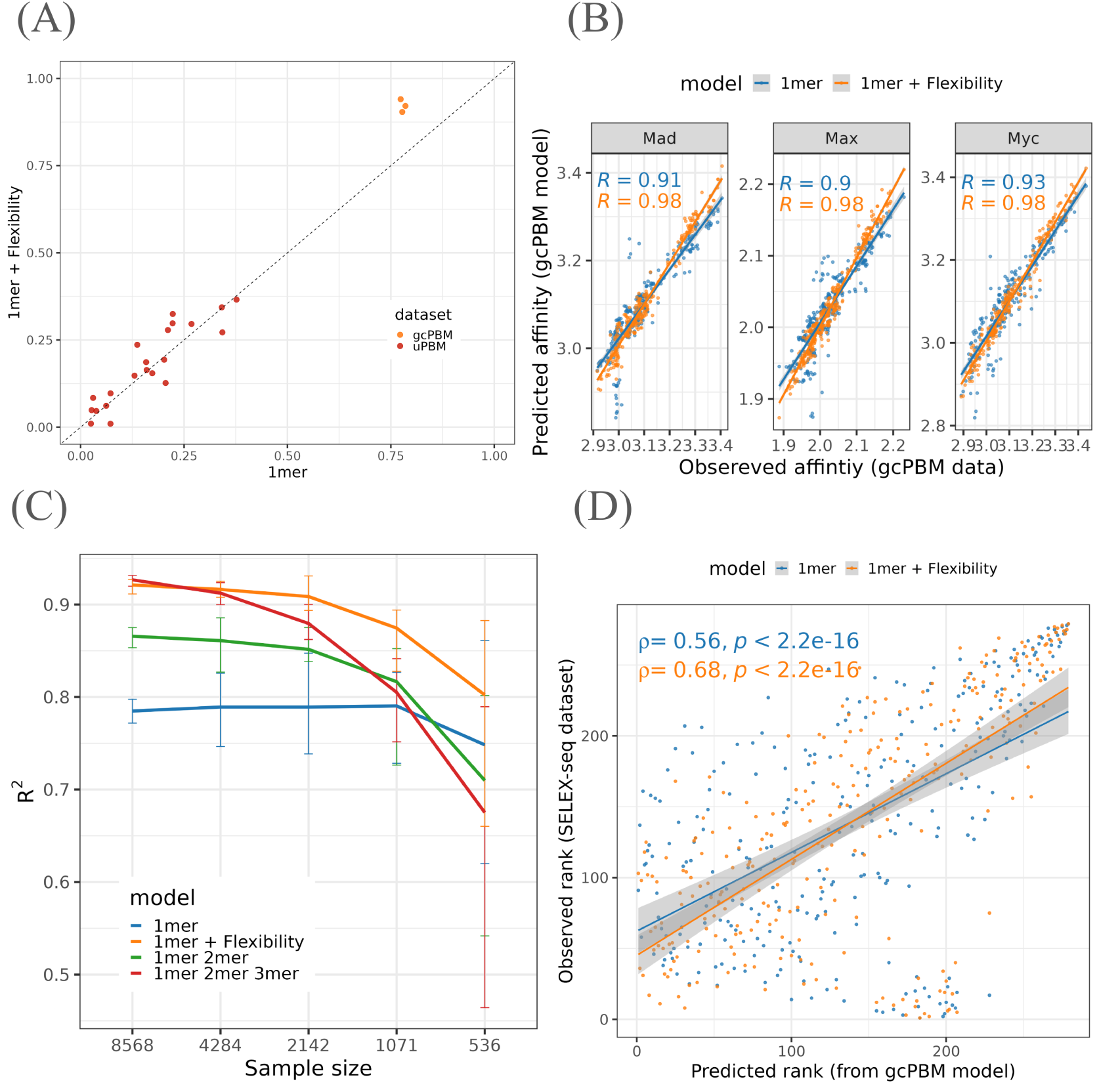
DNA Flexibility Enhances TF Binding Prediction Across Experimental Platforms. **(A)** Comparison of R² values from sequence-only and flexibility-augmented models using universal PBM (uPBM) and genomic-context PBM (gcPBM) datasets. **(B)** Scatter plots of predicted versus observed binding affinities for three bHLH TFs (Mad, Max, c-Myc) showing improved fit with flexibility-augmented models. **(C)** Effect of training dataset size on model performance comparing sequence-only, flexibility-augmented, and higher-order sequence models. **(D)** Cross-platform validation, Spearman’s rank correlation of predicted versus observed TF binding ranks for Max using gcPBM-trained models tested on SELEX-Seq data, demonstrating improved prediction with flexibility features.

Despite the results, we recognized inherent limitations of uPBM datasets. Short probe lengths limit the contribution from flanking nucleotide information, and variability in the positioning of core motifs introduces considerable noise (Ruan and Stormo 2017). In contrast, genomic-context PBM (gcPBM) experiments circumvent these issues by improving design principle, i.e., utilizing longer DNA probes with core 8-bp binding motifs centrally aligned, thus providing higher-quality data and a more accurate representation of TF-binding sites (TFBS) (Bulyk 2007). Hence, we next focused on gcPBM datasets for three basic helix-loop-helix (bHLH) TFs: Max, Mad, and c-Myc (Zhou et al. 2015). Flexibility-augmented models delivered substantial performance improvements for these TFs, reaching exceptionally high R² values (∼0.98) and exhibiting excellent agreement between predicted and experimentally measured affinities (**Figure 3A**). Crucially, predictive performance on gcPBM datasets notably exceeded results obtained from uPBM or HT-SELEX datasets. Furthermore, for all three TFs analyzed using gcPBM data, flexibility-augmented models consistently yielded higher Pearson correlation coefficients between experimental and predicted affinities compared to sequence-only models(**Figure 3B**).

Given that incorporating flexibility descriptors inherently increases model complexity (additional features per nucleotide relative to simpler 1-mer models), we systematically assessed the impact of both dataset size, and model complexity on predictive performance. Using progressively smaller subsets of gcPBM data, flexibility-augmented models consistently maintained superior accuracy compared to sequence-only models (**Figure 3C**). Additionally, we compared flexibility-augmented models against higher-order sequence models with comparable or even greater complexity (1-mer+2-mer, 1-mer+2-mer+3-mer). While higher-order sequence models achieved performance comparable to flexibility-augmented models when trained on the complete dataset, their performance drastically declined with reduced dataset sizes (**Figure 3C**). In contrast, the robustness of flexibility-augmented models highlights the intrinsic biological relevance and utility of DNA flexibility descriptors, particularly under conditions of limited training data (Zhou et al. 2015).

Lastly, to rigorously validate model generalization and robustness against experimental platform-specific noise, we conducted cross-platform model testing. We trained both sequence-only (1-mer), and flexibility-augmented models for the human TF Max using gcPBM data, and subsequently tested these models’ predictive performance on an independent SELEX-Seq dataset (Zhou et al. 2015). Since gcPBM assays measure binding intensity whereas SELEX-Seq yields relative affinity rankings, a direct comparison of R² values was inappropriate (Zhou et al. 2015). Instead, we quantified predictive accuracy using Spearman’s rank correlation (rho) between actual and predicted ranks of Max-binding 10-mer sequences. As depicted in **Figure 3D**, the inclusion of DNA flexibility descriptors substantially improved predictive accuracy, improving rho from 0.56 (sequence-only) to 0.68 (sequence + flexibility). This enhancement confirms that DNA flexibility features significantly improve modeling of TF-binding affinity across experimental platforms.

### DNA Flexibility-Augmented Models Elucidate TF Family-Specific Indirect Readout Mechanisms

DNA flexibility, derived from sequence-dependent properties measured through *in-vivo* experiments, significantly influences TF–DNA binding specificity across diverse organisms. Despite this, determining the precise contribution of flexibility features at individual nucleotide positions within transcription factor binding sites (TFBS) remains underexplored—particularly for mammalian TFs in a family-specific context. Here, we exploited high-quality mammalian

HT-SELEX datasets coupled with flexibility-augmented models to quantify indirect readout mechanisms at base-pair resolution, even in the absence of experimentally solved co-crystal structures (Zhou et al. 2015).

To dissect nucleotide-specific contributions, we defined a quantitative strategy similar to (Yang et al. 2017). For each TF, we first established a baseline performance (𝑅^2^) using a sequence-only (1-mer) model. We then recalculated 𝑅^2^ after incorporating flexibility descriptors at each nucleotide position 𝑖 (1-mer + Flexibility_𝑖_). The performance difference at position 𝑖 was expressed as Δ𝑅_𝑖_^2^ = 𝑅^2^(1-mer + Flexibility_𝑖_) − 𝑅^2^(1-mer), and normalized by the sequence-only model 𝑅^2^ to yield the ratio Δ𝑅_𝑖_^2^/𝑅^2^(1-mer). A positive ratio thus indicated enhanced predictive accuracy attributed directly to flexibility features at a given nucleotide, reflecting its contribution to the indirect readout mechanism along the TFBS.

We applied this approach to systematically assess bendability readout contributions among diverse TF families, and visualized these insights as heatmaps. Rows correspond to TFs, while columns represent nucleotide positions within their respective binding sites (*Supplementary Figure 3A*). To further delineate the role of specific DNA flexibility descriptors—DNaseI sensitivity, nucleosomal positioning preference (NPP), twist-dispersion, and twist-roll-x displacement (trx)—we expanded our analysis to individually examine each flexibility feature’s impact across nucleotide positions.

Our analysis revealed distinct DNA bendability profiles employed by homeodomain TFs (**Figure 4A**). Hierarchical clustering based on these flexibility patterns clearly grouped TAAT-recognizing homeodomain TFs into distinct clusters (*Supplementary* Figure 3A). This clustering based on the relative contribution of DNA flexibity at each nucleotides suggested a complex interplay between sequence recognition preferences and flexibility-mediated indirect readout mechanisms. Notably, for TAAT-recognizing TFs, nucleotides at the 3 side of the binding site consistently exhibited the strongest flexibility contributions (**Figure 4A**). Axial flexibility (DNaseI and NPP) was particularly influential at the 3 -flank, whereas torsional flexibility descriptors (twist-dispersion and trx) contributed at both the 5 and 3 flanking regions (**Figure 4B**). A preliminary analysis for bZIP TFs similarly suggested significant indirect readout contributions obtaining primarily from 3 -flanking nucleotides of their core binding sites (*Supplementary figure 3A*). In contrast, for C2H2 factors these signal are prevalent in the 5’ flanking sites (*Supplementary figure 3A*). Positional importance of DNA flexibility features for other TF families are illustrated in *Supplementary figure 3A-C*.

**Figure 4.**
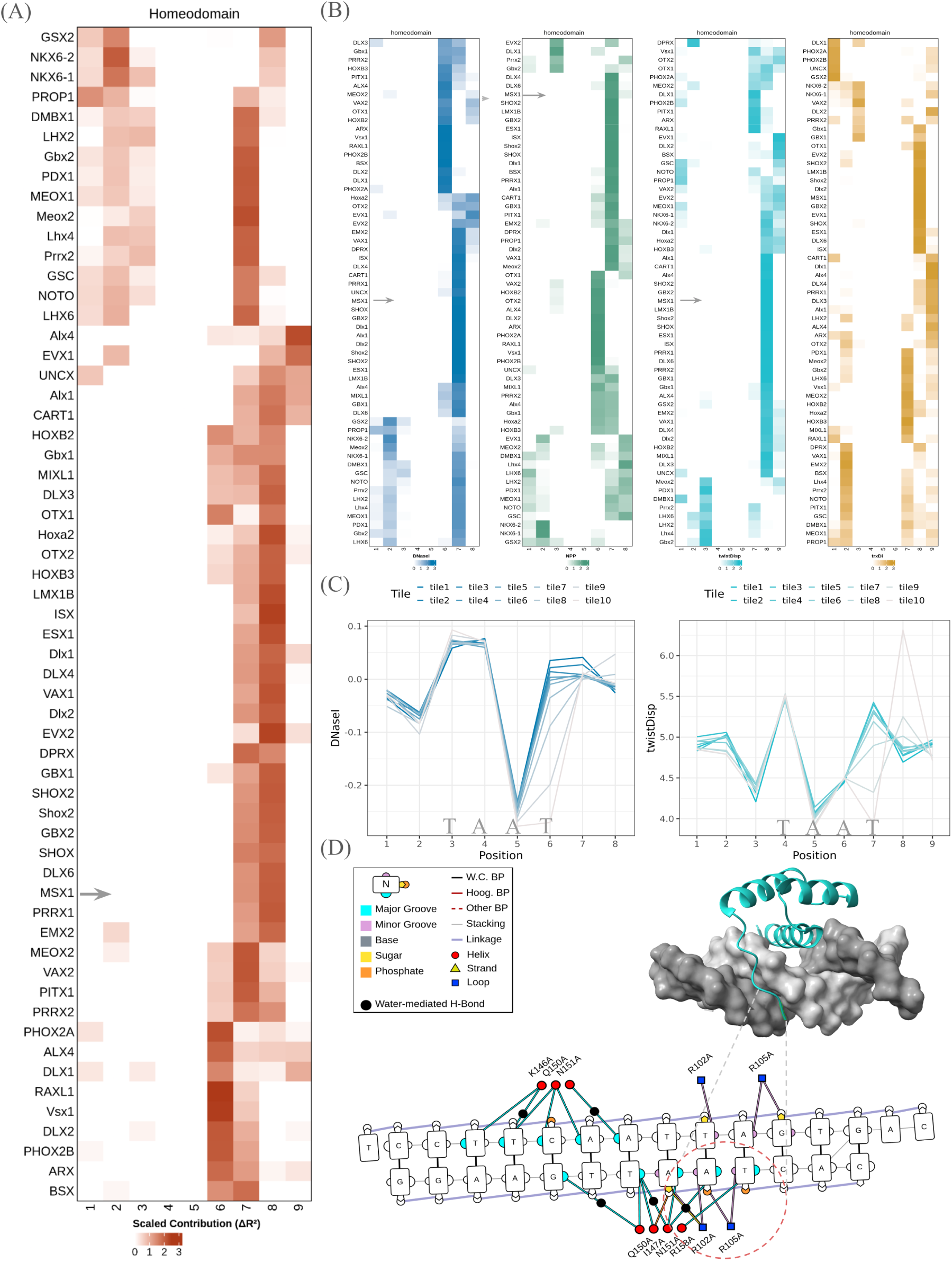
Position-Dependent Contributions of DNA Flexibility to TF Binding Specificity Revealed by Machine Learning with Feature Selection. **(A)** Heatmap showing the impact of adding DNA flexibility features at individual nucleotide positions to a sequence-only model for homeodomain TFs recognizing the TAAT motif. Each row represents a TF, and columns correspond to nucleotide positions within the binding site. **(B)** Heatmaps illustrating the contributions of individual DNA flexibility features at each nucleotide position when combined with sequence-only models for the same set of TFs. **(C)** Average DNA flexibility profiles for DNaseI sensitivity (*left*) and twist-dispersion (*right*) across ten equal-sized bins of MSX1 binding sites. Binding sites were stratified based on normalized MSX1 affinity from HT-SELEX data, with Tile 1 representing the highest affinity sequences and Tile 10 the lowest. **(D)** DNA–protein contact map derived from the co-crystal structure of the MLX1 homeodomain bound to DNA (inset: MLX1 in cyan, DNA in grey). The red dotted circle highlights arginine residues inserted into the minor groove of the consensus TAAT motif.

To validate the findings from feature importance analysis, we examined MSX1, a homeodomain TF. MSX1 binding sites were ranked (M-words from HT-SELEX) into ten affinity-stratified bins, and compared DNA bendability profiles at each nucleotide position. As illustrated in **Figure 4C** (left panel), sequences with highest affinity (*tile 1*) possessed higher DNaseI bendability scores at their 3 -flanking positions of core ‘TAAT’ motif. Given the DNaseI bendability ranges from -0.281 (AAT/ATT trinucleotides) to 0.194 (TCA/TGA), the elevated positive scores indicated a highly flexible trinucleotide environment. This trend aligned closely with the relative flexibility contributions observed in feature importance heatmaps (**Figure 4B**). Likewise, twist-dispersion profiles exhibited a similar localized flexibility at the 3 flank of high-affinity MSX1 sites (**Figure 4D**, right panel). These findings further suggests the affinity variation across MSX1 binding sequences can be attributed to the differences observed in DNA flexibility at the 3 flanks of TAAT motifs.

Finally, to corroborate the observed patterns in the heatmap and flexibility profiles with direct structural evidence, we examined the co-crystal structure of MSX1 DNA complex (Hovde, Abate-Shen, and Geiger 2001) (PDB ID: **1IG7**; **Figure 4D**). The residue interaction map from the structure revealed the MSX1 N-terminal domain engaging the major groove at the TAAT motif by Asn51 and Gln50, inducing narrowing of the adjacent minor groove. Consequently, arginine residues within the MSX1 N-terminal tail interact closely with the narrowed minor groove, potentially contributing to its widening (**Figure 4D** bottom panel, and *Supplementary Figure 3B (C-F)*). Since, these crucial structural dynamics are conducive of MSX1–DNA interactions and facilitate precise recognition, high-affinity sequences likely accommodate essential DNA shape changes via increased flexibility at the 3 -flanking nucleotides as observed in **Figure 4A-C**. Therefore, the structural data align closely with the significant contributions of bendability, particularly the axial bendability, observed at the 3 -flanking region in flexibility-derived feature importance heatmaps (**Figures 4A–B**).

Collectively, these affinity-driven observations are coherent with both our DNA flexibility analyses and structural insights, providing a comprehensive mechanistic view of TF–DNA recognition driven by indirect readout mechanisms.

### DNA Flexibility-Augmented Models Improve Predictive Performance for TF Binding Specificity In Vivo

To evaluate the biological relevance of DNA flexibility, we investigated its impact on predicting transcription factor (TF) binding *in-vivo*. We hypothesized that DNA flexibility could distinguish functional, and occupied TF binding sites (TFBS), from unbound genomic motifs across diverse cellular contexts. To test this, we implemented a classification framework adapted from prior work (Mathelier et al. 2016; Barozzi et al. 2014; Dey et al. 2025), integrating publicly available ENCODE datasets for TF occupancy (ChIP-seq) and chromatin accessibility (DNase-seq). We compared flexibility-augmented models (1-mer + flexibility) against sequence-only baseline models (1-mer), using Matthews Correlation Coefficient (MCC) and Area Under the ROC Curve (AUC) to quantify performance gains attributable to DNA flexibility.

This framework was applied to nearly 900 ENCODE TF ChIP-seq datasets from multiple cell lines. Following the classification approach, we applied a quality control filter, retaining only those datasets where flexibility-augmented models matched or exceeded baseline performance, consistent with expectations that informative features should not degrade model accuracy (Yang et al. 2017). This yielded 307 high-confidence datasets for further analysis.

Across these 307 valid datasets, incorporating DNA flexibility information consistently improved classification performance, with 88.2% showing an increased MCC. As illustrated in **Figure 5A** (also check *Supplementary figure 4A*), where each point represents a dataset, the general shift above the diagonal line signifies widespread predictive gains with the addition of the DNA flexibility descriptors. Given that many TFs were profiled across multiple cell lines, we observed both differences and similarities in performance gains across these cellular contexts (*Supplementary* Figure 4B), demonstrating the robustness of DNA flexibility features. Nevertheless, the trend of improvement retained when datasets were grouped by the 180 unique TFs across different cell-lines, and median MCC improvements were compared between the two models (**Figure 5B**).

**Figure 5.**
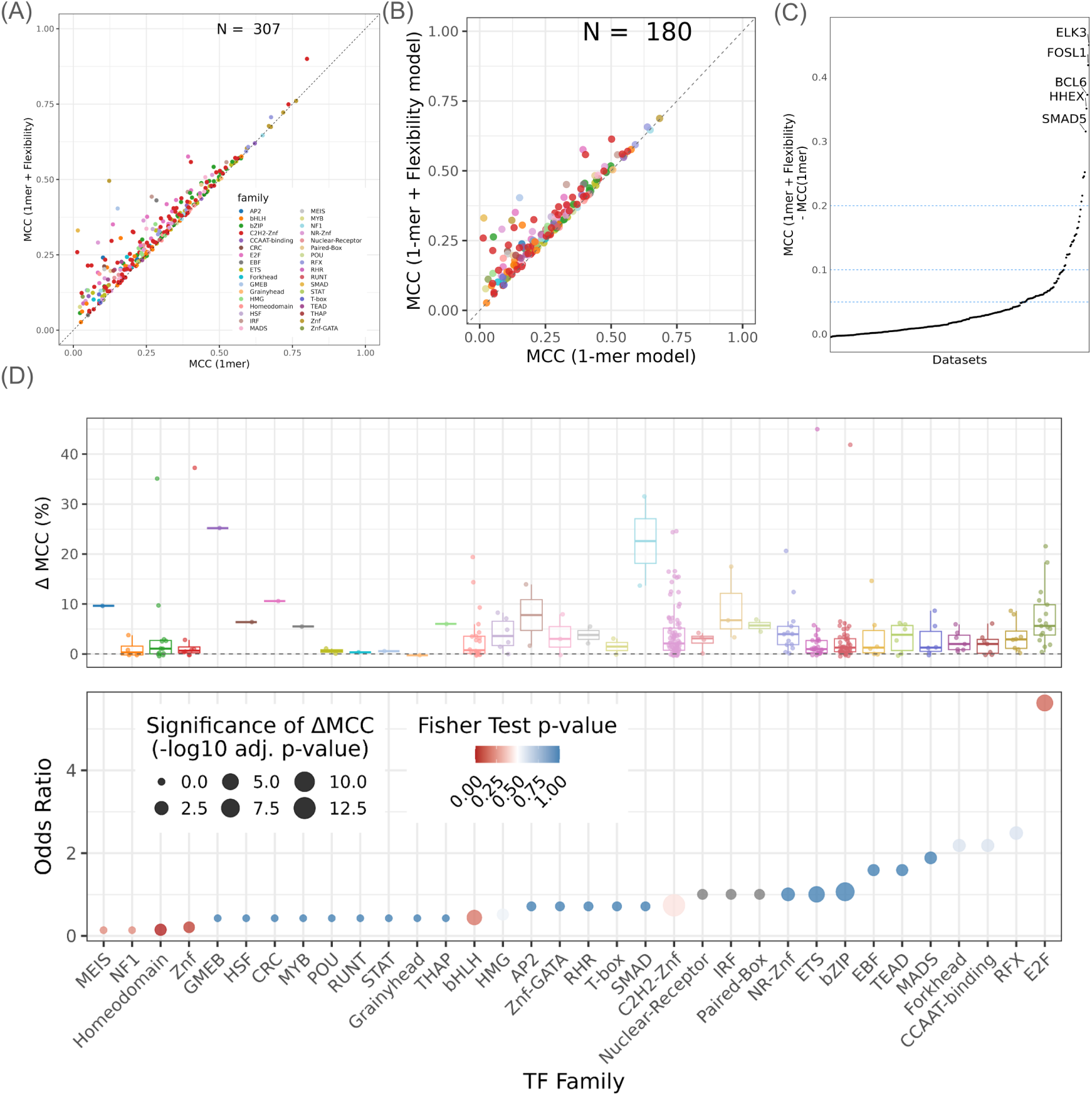
DNA Flexibility-Augmented Models Enhance In Vivo TF Binding Site Prediction Across Diverse TF Families and Cell Types. **(A)** Scatter plot comparing Matthews Correlation Coefficient (MCC) for flexibility-augmented (1-mer + flexibility) versus sequence-only (1-mer) models across 307 high-confidence ENCODE ChIP-seq datasets. Each point represents a dataset; points above the diagonal indicate improved performance with flexibility features. **(B)** Comparison of median MCC values for 178 unique TFs, grouped across cell lines, between flexibility-augmented and sequence-only models. **(C)** Ranked dotplot of MCC improvement (ΔMCC) for each of the 307 datasets. The dashed blue lines denote thresholds of 5%, 10%, and 20% MCC improvement. Selected datasets with MCC improvement exceeding 20% are labeled, highlighting TFs such as ELK3, FOSL1, BCL6, HHEX, and SMAD5. **(D)** *Top:* Distribution of median MCC improvement (ΔMCC %) for each TF family, illustrating family-specific variability due to addition of flexibility features. *Bottom:* Bubble plot showing odds ratios for flexibility-mediated performance improvement by TF family. Bubble size corresponds to median of ΔMCC for each TF-family(also medians in boxplot on top), while color reflects false discovery rate (FDR) corrected p-values (Mann-Whitney U test p-values obtained by comparing distribution of MCC for 1mer vs 1mer+Flexibility models). TF families such as E2F, NR-Znf, Forkhead, and bHLH show odds-ratio more than one, with statistically significant difference in performance gain. Statistical significance of performance differences between two models for each TF families, highlighting those with OR > 1 and significant p-values (Mann-Whitney U test, FDR-corrected).

We quantified MCC gains per dataset, finding that approximately 75% of datasets showed modest improvements (<5%), 15% had moderate gains (5–10%), and 7% exhibited substantial increases (10–20%) (**Figure 5C**), while only ten datasets—including BCL6, E2F3, ELK3, FOSL1, and GMEB2—obtained MCC improvements exceeding by 20%. Notably, TFs demonstrating modest to substantial improvements spanned diverse families and recognition mechanisms, aligning with our hypothesis that DNA bendability broadly contributes to TF recognition across distinct TF families.

Lastly, in order to assess the family-specific relevance of DNA flexibility, we grouped TFs by family and compared median MCC gains. **Figure 5D (top)** reveals variable performance improvements among the TF families. For rigorous statistical comparison of performance improvements while accounting for biased sampling of TFs per family in the current dataset, we calculated an enrichment odds ratio (OR). As shown in **Figure 5D** (*bottom*), The E2F family exhibited the highest OR, indicating a five fold increased likelihood of improvement by DNA flexibility features. Moreover, six other TF families—including EBF, TEAD, MADS, Forkhead, CCAAT-binding, and RFX showed ORs greater than one. Nonetheless, we also observed statistically significant median MCC differences (Mann-Whitney U test, FDR-corrected p-values) from flexibility augmented models compared to the baseline models. These results further undeline that DNA bendability is a relevant determinant of *in-vivo* functional TFBS prediction, with broader physiological context, albeit with varying degrees of impact across TF-families.

## Discussions

Precise transcription factor (TF) binding to cognate DNA sequences is central to eukaryotic gene regulation. Despite decades of structural and biochemical research elucidating protein– DNA interfaces, the comprehensive principles governing TF specificity and the cis-regulatory code remain elusive (Boer and Taipale 2024; Kim and Wysocka 2023). It is now widely recognized that TF–DNA recognition arises from a multifaceted interplay among core motif sequences, flanking DNA context, nucleosome positioning, and chromatin architecture in vivo (Grand et al. 2024). Recently, increasing focus has shifted toward higher-order DNA structural features—particularly shape and flexibility—and their combinatorial influence on binding specificity (Chiu et al. 2022; Liu et al. 2025; Li, Chiu, and Rohs 2024; Back and Walther 2023; Mitra et al. 2023). Notably, while current DNA shape-based models acknowledge the relevance of flexibility in DNA–TF interactions, they often exclude this critical parameter in predicting TF binding sites, possibly due to experimental limitations in validating flexibility at very short oligonucleotide scales (Li et al. 2017).

Our systematic reanalysis of extensive TF–DNA binding datasets underscores the pivotal role of DNA flexibility, supported by experimental and computational evidence, in enhancing the accuracy of predictive models of TF–DNA interactions. These findings substantiate the concept that local DNA conformational landscapes modulate TF binding in ways that extend beyond direct base-specific contacts (Rohs et al. 2010). Prior studies have reported the importance of DNA flexibility for a limited range of TFs (Yella et al. 2018; Dey et al. 2025, 2023; Murthy et al. 2024; Sarkar et al. 2021); our work generalizes these observations across a broad spectrum of TF families, bridging a critical gap in understanding how sequence-encoded DNA structural dynamics govern specificity. Consistently, models incorporating di- and trinucleotide flexibility yield markedly improved predictive performance across multiple TF classes, emphasizing the nuanced and complementary contributions of axial, torsional, and stretching flexibility to indirect readout mechanisms.

A central insight from our analysis is inherent ability of DNA to adopt structural conformations favorable for TF binding, complementing base-specific recognition (Rohs et al. 2010). This synergy arises because sequence-encoded conformational plasticity facilitates or restricts local shape transitions essential for stable TF–DNA complexes, particularly at the major and minor groove interfaces (Ghoshdastidar and Bansal 2022). By decomposing TF binding site flexibility into axial, torsional, and stretching components, we gain mechanistic clarity on these interactions. Supporting this, principal component analyses reveal that integrating DNA flexibility with sequence information captures a greater proportion of variance in TF–DNA binding data, resulting in more distinct clustering of TFs by family and highlighting shared yet diverse recognition strategies. Notably, families such as bHLH and bZIP exhibit unique “flexibility signatures” that sharply contrast with homeodomains, reflecting fundamentally different DNA recognition modalities and sequence motif architectures.

Crucially, these insights underscore the intricate interdependencies among DNA sequence, shape, and flexibility in directing TF binding. DNA flexibility functions as a key mediator between the primary sequence and its resulting shape. Remarkably, even a perfect cognate sequence can be driven into suboptimal DNA conformations by “flexibility signatures” embedded in flanking sequences (Ghoshdastidar and Bansal 2022). This results in TF binding sites sampling a broad “shape space” of sequence-encoded conformations, with only a subset exhibiting high-affinity binding due to favorable flexibility patterns. Conceptually, sequence-only (1-mer) models capture the direct, base-specific interactions critical for canonical hydrogen bonds and van der Waals forces, while flexibility descriptors quantify the DNA’s propensity to adopt geometries compatible with these contacts. The integration of these complementary layers consistently enhances TF–DNA specificity models across families, yielding a more accurate and biologically faithful depiction of the underlying binding energy landscape.

The critical role of DNA flexibility is further highlighted by significant gains in predictive accuracy observed with flexibility-augmented models compared to sequence-only baselines in vitro. Although some TF families show more modest improvements—potentially due to limited dataset representation—the majority exhibit notable enhancement, supporting the widespread relevance of flexibility-dependent indirect readout. In agreement, ridge regression analyses emphasize the particular importance of di- and trinucleotide flexibility features in modeling the binding landscapes of bHLH and homeodomain TFs, reinforcing similar conclusions drawn from DNA shape perspectives (Yang et al. 2017).

Moreover, these advantages transcend experimental platforms and species boundaries. Consistent improvements in predictive performance (𝑅^2^) are observed across diverse datasets, including Drosophila Hox SELEX-Seq, mouse universal protein binding microarrays (uPBM), and human genomic-context PBM (gcPBM) datasets. This cross-platform reproducibility strongly suggests that local DNA bendability is a biologically conserved and intrinsic determinant of TF-mediated DNA deformation. Supporting this, flexibility-augmented models trained on one dataset (e.g., Max TF from gcPBM) successfully predict binding affinities in independent assays (e.g., SELEX-Seq), confirming that sequence-encoded bendability captures a fundamental dimension of TF–DNA energetics. Importantly, compared to higher-order sequence features (2-mer, 3-mer), DNA flexibility features provide a more cost-effective solution with comparable or superior generalization and lower dimensionality—a finding that aligns with prior observations on DNA shape features (Zhou et al. 2015).

Finally, dissecting axial, torsional, and stretching flexibility at individual nucleotide positions reveals position-specific “flexibility hotspots” within TF binding sites and their flanking regions. Homeodomains recognizing TAAT motifs show pronounced flexibility contributions at the 3 flanks, whereas bZIP factors exploit flanking flexibility extensively, likely to accommodate their characteristic “DNA clamp” binding architecture. This DNA flexibility-augmented framework transcends anecdotal structural observations by systematically quantifying bendability contributions across multiple TF families, providing a unified mechanistic understanding of indirect readout in TF–DNA recognition. Unlike many TFs, homeodomains are known to directly recognize flanking sequences, a property clearly reflected in our position-specific bendability heatmaps. For example, high-affinity TAAT motifs recognized by MSX1 are flanked by sequences with structural features favorable for binding. MSX1 preferentially binds to highly flexible 3 flanks that facilitate narrower minor groove conformations, recognized by arginine residues, followed by key major groove interactions mediated by Asn51 and Gln50 at the core TAATTG motif (Rohs et al. 2010).

The biological relevance of DNA flexibility is further corroborated in vivo. Applying a classification framework integrating ENCODE ChIP-seq and DNase-seq data, we distinguished functional TF binding sites from unbound genomic motifs. After quality filtering of over 900 datasets, 307 high-confidence datasets demonstrated that incorporating DNA flexibility improved classification performance in 88.2% of cases. Although the magnitude of improvement varied, TFs such as BCL6, E2F3, and FOSL1 exhibited substantial gains. These predictive enhancements across diverse TFs and cellular contexts underscore DNA flexibility as a biologically meaningful signal in in-vivo binding site recognition.

Family-specific analyses reinforced these findings. Calculating enrichment odds ratios to adjust for family size revealed that TF families such as E2F had a fivefold increased likelihood of improved model performance when incorporating flexibility features. Several other families— C2H2 zinc finger, Nuclear Receptor, bZIP, ETS, Forkhead, RFX, and NR-Znf—also showed statistically significant improvements in MCC. These results align with previous reports highlighting flexibility’s critical role in prevalent TF families in eukaryotes (Yella et al. 2018). Together, our data extend in vitro conclusions into the in vivo environment, firmly establishing DNA bendability as a broadly relevant determinant of functional TF binding site recognition, with family-specific mechanistic implications.

## Conclusion

Our findings collectively underscore the essential role of DNA flexibility in transcription factor (TF)–DNA recognition. By systematically integrating di- and trinucleotide bendability scales with sequence-based (1-mer) motifs, we provide compelling evidence that indirect readout mechanisms broadly contribute to binding specificity across diverse TF families. This expanded perspective—where DNA mechanics and sequence together define the binding landscape, opens new avenues for improving predictive models of gene regulation. Importantly, it highlights flanking DNA not merely as passive sequence context but as an active participant in TF binding events.

## Methodology

### DNA-Protein Binding Datasets and Preprocessing

We utilized multiple *in-vitro* transcription factor (TF)–DNA binding specificity datasets generated from distinct experimental platforms. The primary dataset comprised HT-SELEX-derived DNA-binding affinities for 215 mammalian TFs spanning 27 protein families (*ENA accession PRJEB14744*) (Yang et al. 2017). Pre-processed TF M-word datasets with affinity scores were obtained from the Rohs Lab website. Additionally, SELEX-Seq datasets for Drosophila Hox TFs in complex with the Exd cofactor were retrieved from the Gene Expression Omnibus (GEO accession GSE65073) (Abe et al. 2015). For these datasets, 16-bp SELEX-Seq sequences were scanned for Hox binding sites using the consensus motif “TGAYNNAY,” followed by filtering for motifs centrally located within sequences, which were then trimmed to 14 bp for analysis (Zhou et al. 2015).

Genomic-context protein binding microarray (gcPBM) data for dimeric bHLH TFs Max/Max, Mad1/Max, and c-Myc/Max were obtained from GEO (accession GSE59845) (Zhou et al. 2015). These datasets consist of 36-mer probes embedded in native genomic sequences, with centrally aligned E-box motifs, allowing direct analysis without additional filtering. Cross-platform prediction of Max binding affinity was conducted by processing gcPBM data as described in Zhou et al (Zhou et al. 2015). Universal PBM (uPBM) data for 66 mouse TFs, representing various protein families, were sourced from GEO accession *GSE42864* (Weirauch et al. 2013). Normalized signal intensities were used for analysis. Due to the unaligned nature of uPBM probes, both unaligned and aligned trimmed sequences were considered following protocols from Zhou et al (Zhou et al. 2015).

### DNA Conformational Flexibility Predictions

DNA is a dynamic molecule exhibiting sequence-dependent flexibility, shape variations, groove geometry, and electrostatic properties. Protein binding can induce conformational changes including bending and twisting along the DNA. DNA flexibility is decomposed into axial (bending), torsional (twisting), and stretching components (Vanaja et al. 2021; Dey et al. 2025).

Axial flexibility was quantified using dinucleotide and trinucleotide models. Two trinucleotide scales were employed: DNaseI sensitivity (Brukner et al. 1995), derived from *in-vivo* DNaseI cleavage data reflecting bending propensity towards the major groove (higher values indicate greater flexibility); and the nucleosome positioning preference (NPP) (Satchwell, Drew, and Travers 1986), based on histone-bound DNA sequence analysis in chicken erythrocyte nucleosomes (lower values indicate higher flexibility) (Yella, Kumar, and Bansal 2014).

Torsional flexibility was assessed via twist dispersion (Olson et al. 1998; Beno et al. 2011; Inga et al. 2002), measuring variance in helical twist angles at dinucleotide steps from DNA-protein crystal structures (higher values indicate higher twisting flexibility), and twist-roll-x displacement (trxDi), which captures intrinsic dinucleotide conformational variability based on the equilibrium between B-I and B-II sugar-phosphate backbone states (Heddi et al. 2010).

Stretching flexibility, related to DNA compressibility, was quantified from molecular dynamics-derived stretch moduli of dinucleotide steps, with lower modulus values indicating increased flexibility (Marin-Gonzalez et al. 2019).

### Encoding DNA Flexibility Features of TFBS

For each of the datasets from SELEX, gcPBM and other experiments, the sequences were one-hot encoded at the nucleotide level. DNA flexibility features were incorporated using sliding windows: trinucleotide-based DNaseI and NPP scales, and dinucleotide-based twist dispersion, trxDi, and stiffness models implemented in the python package DNAflexpy.

### Principal Component Analysis (PCA)

For each TF, the top affinity sequence was encoded using 1-mer features alone, then complemented by the four flexibility descriptors (DNaseI, NPP, twist dispersion, and trxDi) representing axial and rotational flexibility. PCA was performed in R with prcomp package (R Core Team et al. 2013), separately on sequence-only and sequence-plus-flexibility encodings. Inter- and intra-family Euclidean distances were calculated from the first two principal components to assess clustering.

Regression models to relate ***k***-mer affinity to sequence and bendability features

To enumerate the relationship of relative binding affinities with DNA sequence and sequence dependent bendability, we employed L2-regularised multiple linear regression (L2-MLR). For an affinity vector 𝑦

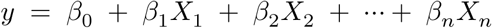

𝑦 contains the measured affinities for each *k*-mer, the 𝑋 are features that encode the corresponding DNA sequence, 𝛽_𝑗_are regression coefficients, and 𝛽_0_is an intercept. To prevent overfitting, L2-regularization employs an additional penalty term on the coefficients in the loss function 𝐿(𝛽). Then coefficients are obtained by minimizing

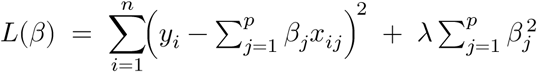

with the regularisation strength 𝜆 chosen by nested cross-validation from a range of 𝜆 ∈ [10^−1^, 10^6^].

The baseline (1-mer) models are mononucleotide representation with one-hot encoded features. For a binding site of length 𝑘, four binary features encode the identity of each nucleotide position, so the design matrix has 4𝑘 features. Bendability-augmented models are labelled 1-mer + Flexibility extend the sequence encoding with five experimentally derived bendability descriptors:

| Models | Representation | Degrees of freedom captured |
| --- | --- | --- |
| DNase I bendability | 3 bp (trinucleotide) | axial bending |
| NPP (nucleosome-positioning preference) | 3 bp | axial bending |
| Twist-dispersion | 2 bp (dinucleotide) | torsional flexibility |
| trx (twist-roll-$x$ displacement) | 2 bp | torsion/roll coupling |
| Stiffness | 2 bp | stretching/compressibility |

DNase I and NPP are assigned by sliding a 3-bp window, yielding 𝑘 − 3 + 1 features. The three dinucleotide scales are assigned by a 2-bp window, yielding 𝑘 − 2 + 1 features each. These feature encoding are performed using DNAflexpy (citation). Concatenating these with the 1-mer matrix results in:

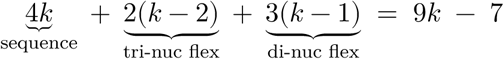

features per *k*-mer. Each flexibility value is 𝑧-score normalized before concatenation so that regression weights remain on comparable scales. Additionally, we also used interaction terms of the each flexibility feature as a second-order representation of nucleotide interdependencies.

In case of gcPBM datasets, 2-mer(dinucleotide) and 3-mer(trinucleotide) were also considered along with the 1-mer. There are (4^2^ = 16) possible dinucleotides ((AA, AC, … , TT), so the feature matrix adds 16(𝑘 − 1), columns, i.e. 16 binary indicators for each of the (𝑘 − 1) overlapping dinucleotide windows. Similarly, for 3-mer (trinucleotide) model– each sliding window of three bases is encoded with (4^3^ = 64, obtaining 64(𝑘 − 2) additional features. In general, an **(N)-mer** model (with (𝑁 ≥ 1)) requires (4^𝑁^ ) indicators for each of the (𝑘−𝑁 +1) windows, combining 4^𝑁^ (𝑘 − 𝑁 + 1) features.

When multiple *N* -mer orders are concatenated, the total dimensionality is the sum of the individual feature counts. For example, a 1-mer + 2-mer model uses 4𝑘 + 16(𝑘−1) = 20𝑘−16 features, or exactly *20* features per nucleotide. A 1-mer + 2-mer + 3-mer stack would similarly require 4𝑘 + 16(𝑘 − 1) + 64(𝑘 − 2). A full equivalences among these encodings and their information content are discussed in detail by Yang *et al*. (Yang et al. 2017).

Model fitting and evaluations were implemented with RidgeCV from *scikit-learn* (Pedregosa et al. 2011). For each TF dataset, a 10-fold outer loop generated train–test splits; within each outer fold, a 5-fold inner loop tuned 𝜆. Final performance on the held-out folds was summarised by 𝑅^2^, mean absolute error (MAE), and standard error (s.e.). Gains in these metrics, relative to the 1-mer baseline, quantify the predictive value of bendability descriptors for each transcription factor.

### Feature Selection for Position-Specific Flexibility Contributions

For each transcription factor (TF), we first fit a baseline ridge-regression model using 1-mer sequence features and recorded its cross-validated coefficient of determination, 𝑅_1_^2^_-mer_.

We then iteratively augmented the feature set by adding the five flexibility scores at a single nucleotide position *i* (DNase I, NPP, twist-dispersion, trx, stiffness), refit the model, and obtained the new goodness-of-fit, 𝑅_1_^2^_-mer+𝐹𝑖_ .

The performance gain at position *i*:

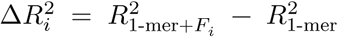

### To compare positions on a common scale, we expressed the gain as a relative contribution

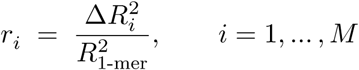

A positive 𝑟_𝑖_indicates that flexibility at nucleotide *i* improves affinity prediction beyond sequence identity alone, highlighting a position-specific bendability-readout. Plotting the 𝑟_𝑖_ profiles across all positions and TFs yields heat maps that reveal how axial (DNase I, NPP), torsional (twist-dispersion, trx), and stretching (stiffness) flexibility contribute to binding specificity at each nucleotides in TFBS.

### Structural Analysis of TF-DNA Complexes

DNA–TF co-crystal structures were analyzed using the DNAproDB web server (Mitra et al. 2025). Structural DNA shape parameters and residue interaction maps were extracted for comparison with flexibility feature importance profiles.

Classification of In Vivo Transcription Factor Binding Sites

To investigate the in vivo relevance of DNA flexibility, we analyzed transcription factor (TF) binding using a classification framework integrating data from the ENCODE project. We curated 940 TF ChIP-seq datasets across five ENCODE cell lines—K562, GM12878, HepG2, MCF-7, and HeLa-S3. TF occupancy regions were defined using Irreproducible Discovery Rate (IDR) thresholded peaks from ENCODE, while matched DNase-seq data delineated open and closed chromatin states. Our analysis focused exclusively on TF binding sites (TFBS) located within open chromatin regions, consistent with established approaches (Dror et al. 2015).

For each TF, the human reference genome (hg38) was scanned using its corresponding Position Weight Matrix (PWM) with the FIMO tool (MEME Suite) (Grant, Bailey, and Noble 2011). Putative motif instances were labeled as positive, occupied TFBS if they met two stringent criteria: (1) the motif plus 50 bp flanking sequences on both sides were fully encompassed within an IDR-thresholded ChIP-seq peak for that TF; (2) the central 100 bp of the ChIP-seq peak was entirely within an open chromatin region, as indicated by DNase-seq peaks.

Genomic intervals meeting these criteria constituted the positive set. Conversely, the **negative set** comprised motif instances located in open chromatin but lacking any overlapping ChIP-seq peak signal within 50 bp upstream or downstream of the motif. DNA sequences for both sets were extracted using BEDTools (Quinlan 2014), forming a high-confidence dataset of bound and unbound motifs for downstream modeling.

We constructed gradient boosting classifiers (XGBoost) using one-hot encoded 1-mer features, and 1-mer combined with DNA flexibility descriptors—DNase I sensitivity, nucleosome positioning preference (NPP), twist dispersion, trx, and stiffness. Classifiers were implemented with the XGBClassifier from XGBoost library in scikit-learn (Pedregosa et al. 2011).

Robust hyperparameter tuning was performed using nested cross-validation on each dataset. Initial modeling utilized 1mer sequence features, and then with 1mer with flexibility descriptors to efficiently explore the parameter space. The outer loop employed stratified 10-fold cross-validation, while an inner 5-fold stratified cross-validation with GridSearchCV optimized hyperparameters over the grid: n_estimators: 50, 100, 200, 300; max_depth: 3, 5, 7, 10; learning_rate: 0.01, 0.1, 0.2, 0.3. Model performance was rigorously evaluated using multiple metrics, including Area Under the ROC Curve (AUC), and Matthews Correlation Coefficient (MCC).

## Data Availability Statement

All source code used for the analysis, scripts to generate the figures, and the underlying processed data supporting the findings of this study are publicly available in a GitHub repository.

