## Supplementary file for "DNA Conformational Flexibility Descriptors Improve Transcription Factor Binding Prediction Across the Protein Families"

Dr. Aditya Kumar[1,*]

2025-08-13

[1] Tezpur University, Molecular Biology and Biotechnology, Tezpur, Assam, 784028, India
[2] Koneru Lakshmaiah Education Foundation, Department of Biotechnology, Vaddeswaram, Guntur, Andhra Pradesh, 522502, India

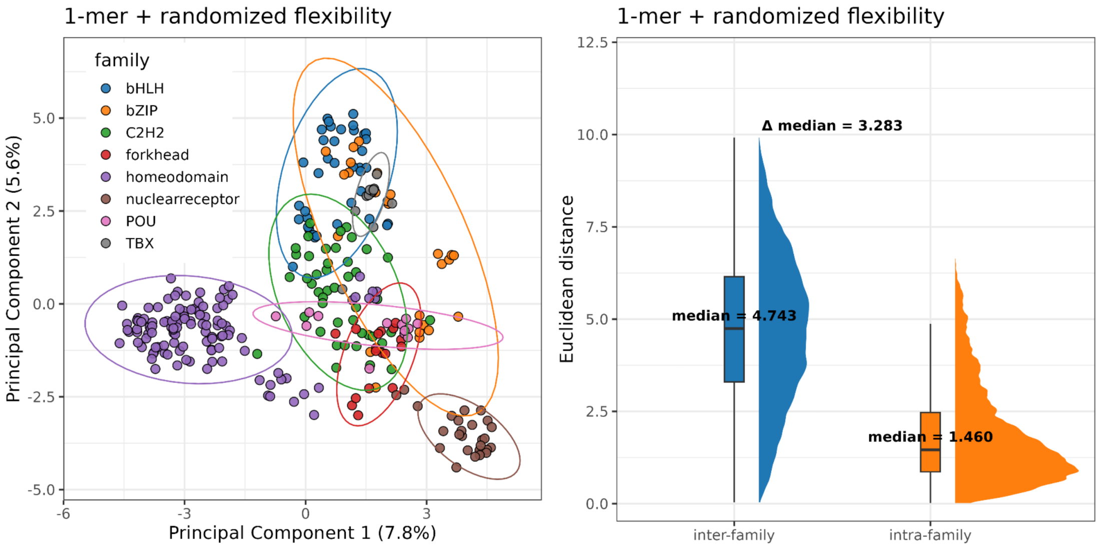
1-mer + randomized flexibility
family
bHLH
bZIP
C2H2
forkhead
homeodomain
nuclearreceptor
POU
TBX
Principal Component 1 (7.8%)
Principal Component 2 (5.6%)
1-mer + randomized flexibility
Δ median = 3.283
median = 4.743
median = 1.460
Euclidean distance
inter-family
intra-family

Figure 1

**Supplementary to Figure 1: Control PCA Using Randomized DNA Flexibility Features Fails to Recapitulate TF Family-Specific Separation:** (*Left*) Principal Component Analysis (PCA) of TF binding sites using 1-mer features combined with randomized flexibility descriptors. Randomized features were generated to match the mean and variance of the original flexibility data. Unlike the real features (see Figure 1A), these surrogate inputs fail to produce meaningful clustering of TFs by family. (*Right*) Distribution of pairwise Euclidean distances between TFs, calculated from the first two principal components. Compared to models using true flexibility features, the randomized model shows reduced separation between inter-family and intra-family TFs. The lower median distance between families illustrates the loss of discriminative power when flexibility descriptors lack biological relevance.

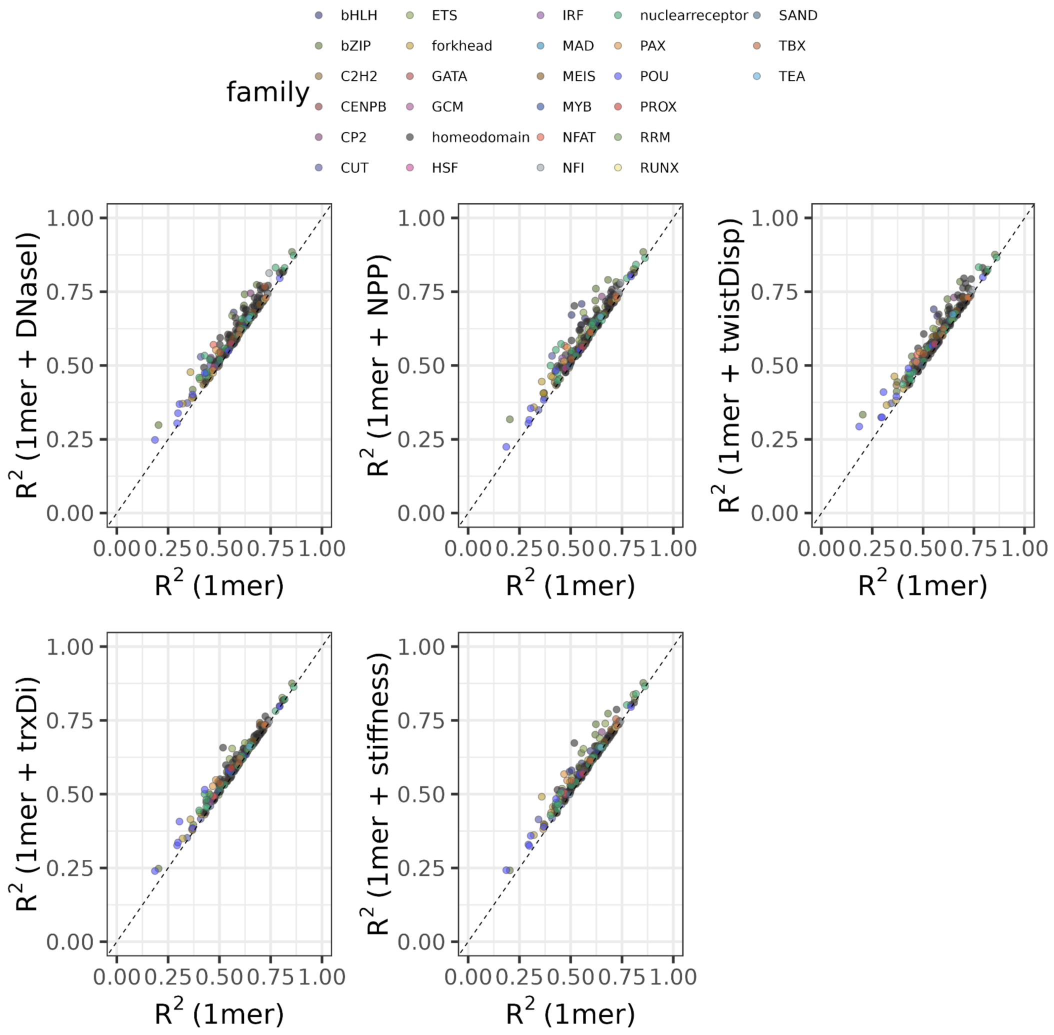
family
bHLH
bZIP
C2H2
CENPB
CP2
CUT
ETS
forkhead
GATA
GCM
homeodomain
HSF
IRF
MAD
MEIS
MYB
NFAT
NFI
nuclearreceptor
PAX
POU
PROX
RRM
RUNX
SAND
TBX
TEA
R^2 (1mer + DNaseI)
R^2 (1mer + NPP)
R^2 (1mer + twistDisp)
R^2 (1mer + trxDi)
R^2 (1mer + stiffness)
R^2 (1mer)
0.00
0.25
0.50
0.75
1.00

Figure 2

**Supplementary Figure 2A:** Dot plots comparing the predictive performance of L2-regularized linear regression models trained on 1-mer features alone versus models trained on 1-mer features combined with individual DNA flexibility features. Each dot represents the mean $R^2$ value obtained from 10-fold cross-validation for a specific transcription factor (TF) dataset. The $R^2$ values on the x-axis correspond to the model trained using only 1-mer features, while the y-axis shows the $R^2$ for models incorporating an additional flexibility feature (DNAseI, NPP, twistDisp, trxDi, or stiffness). Colors indicate different TF families.

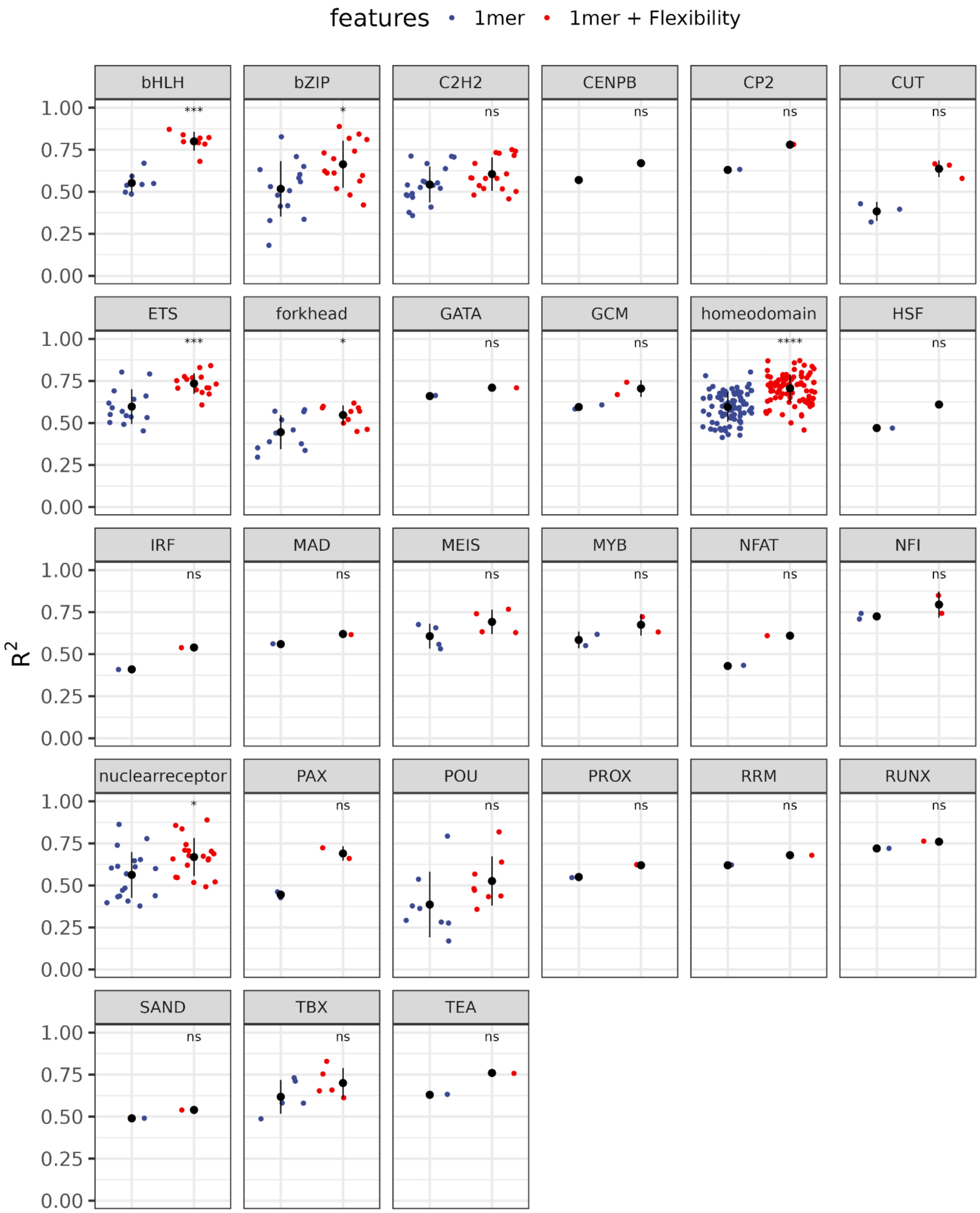

features 1mer 1mer + Flexibility
bHLH
bZIP
C2H2
CENPB
CP2
CUT
ETS
forkhead
GATA
GCM
homeodomain
HSF
IRF
MAD
MEIS
MYB
NFAT
NFI
nuclearreceptor
PAX
POU
PROX
RRM
RUNX
SAND
TBX
TEA
***
*
ns
****
1.00
0.75
0.50
0.25
0.00
R2


Figure 3

**Supplementary Figure 2B:** Distribution of mean $R^2$ values obtained from 10-fold cross-validation for each transcription factor (TF) dataset. Blue and red dots represent models trained with 1-mer features only and 1-mer features combined with DNA flexibility features, respectively. Statistical differences between the two model groups for each TF family were assessed using the Wilcoxon rank-sum test, with significance indicated on the plots. Statistical significance is indicated as: ns ($p > 0.05$), * (p 0.05), ** (p 0.01), *** (p 0.001), **** (p 0.0001).

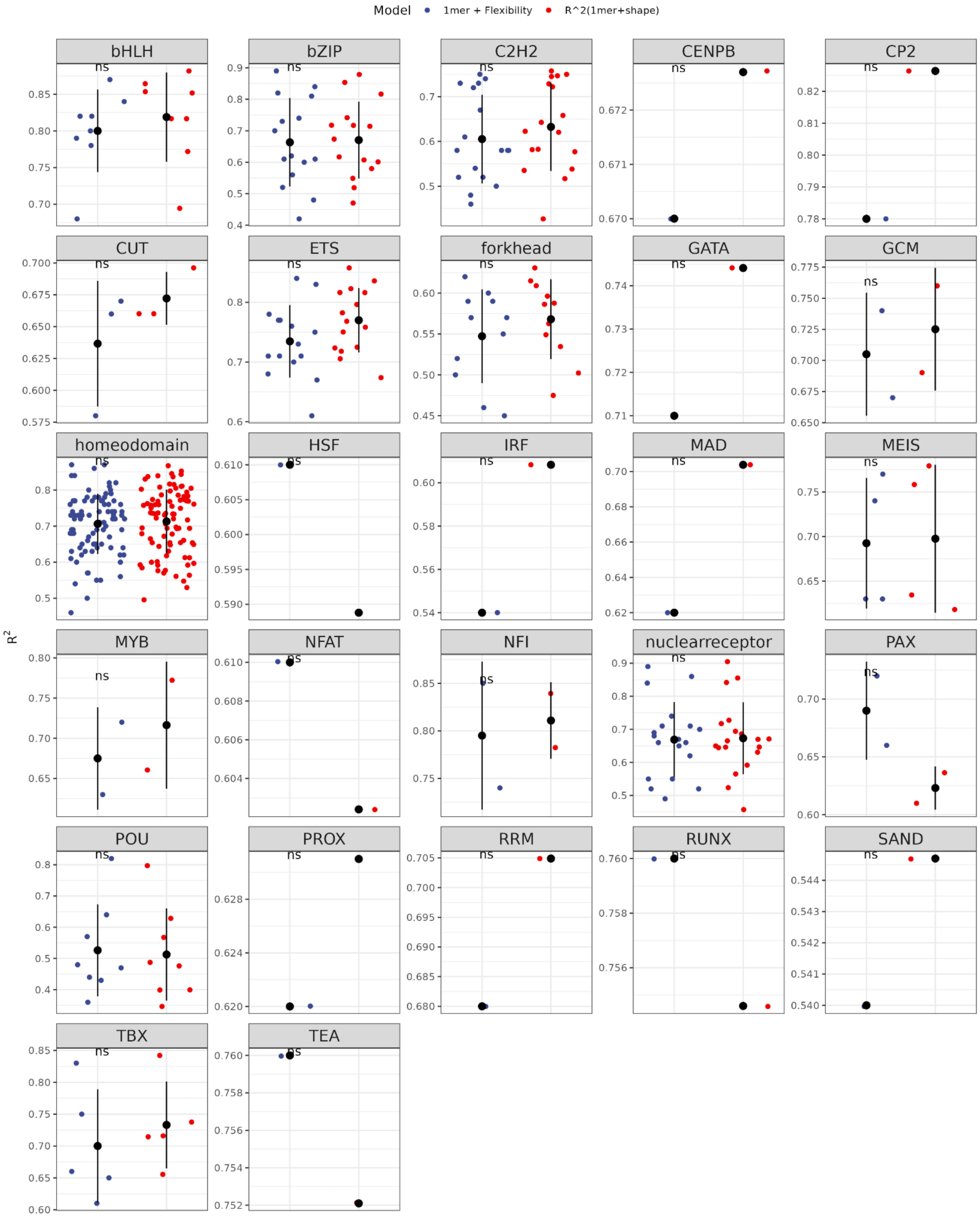

Model
1mer + Flexibility
R^2(1mer+shape)
R²
bHLH
bZIP
C2H2
CENPB
CP2
CUT
ETS
forkhead
GATA
GCM
homeodomain
HSF
IRF
MAD
MEIS
MYB
NFAT
NFI
nuclearreceptor
PAX
POU
PROX
RRM
RUNX
SAND
TBX
TEA
ns


Figure 4

**Supplementary Figure 2C:** Distribution of mean $R^2$ values obtained from 10-fold cross-validation for each transcription factor (TF) dataset. Blue and red dots represent models trained with 1-mer features combined with flexibility features and 1-mer features combined with DNAShape features, respectively. Statistical differences between the two model groups for each TF family were assessed using the Wilcoxon rank-sum test, with significance indicated on the plots. Statistical significance is indicated as: ns ($p > 0.05$), * (p  0.05), ** (p  0.01), *** (p  0.001), **** (p  0.0001).

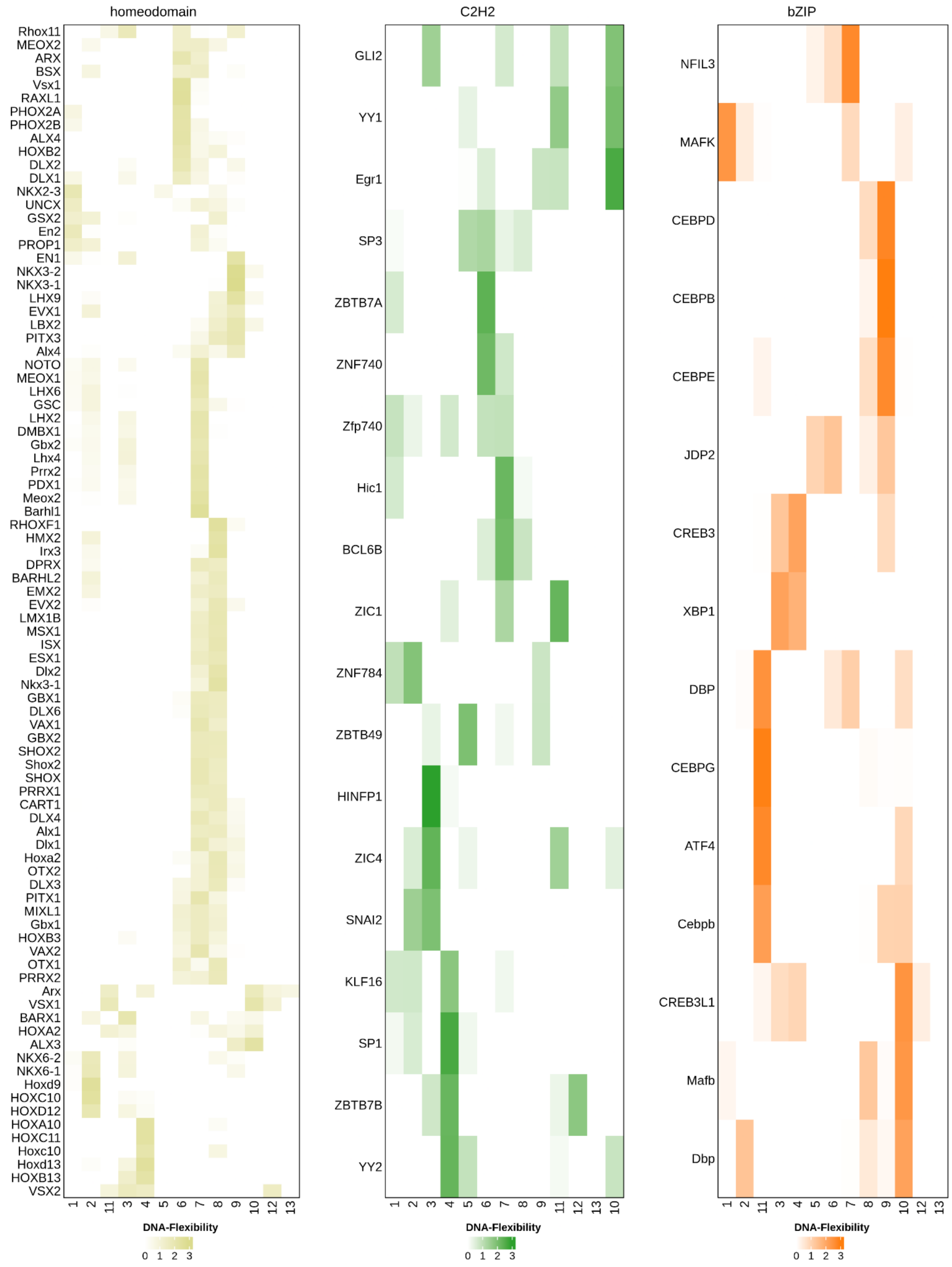
homeodomain
C2H2
bZIP
Rhox11
MEOX2
ARX
BSX
Vsx1
RAXL1
PHOX2A
PHOX2B
ALX4
HOXB2
DLX2
DLX1
NKX2-3
UNCX
GSX2
En2
PROP1
EN1
NKX3-2
NKX3-1
LHX9
EVX1
LBX2
PITX3
Alx4
NOTO
MEOX1
LHX6
GSC
LHX2
DMBX1
Gbx2
Lhx4
Prrx2
PDX1
Meox2
Barhl1
RHOXF1
HMX2
Irx3
DPRX
BARHL2
EMX2
EVX2
LMX1B
MSX1
ISX
ESX1
Dlx2
Nkx3-1
GBX1
DLX6
VAX1
GBX2
SHOX2
Shox2
SHOX
PRRX1
CART1
DLX4
Alx1
Dlx1
Hoxa2
OTX2
DLX3
PITX1
MIXL1
Gbx1
HOXB3
VAX2
OTX1
PRRX2
Arx
VSX1
BARX1
HOXA2
ALX3
NKX6-2
NKX6-1
Hoxd9
HOXC10
HOXD12
HOXA10
HOXC11
Hoxc10
Hoxd13
HOXB13
VSX2
GLI2
YY1
Egr1
SP3
ZBTB7A
ZNF740
Zfp740
Hic1
BCL6B
ZIC1
ZNF784
ZBTB49
HINFP1
ZIC4
SNAI2
KLF16
SP1
ZBTB7B
YY2
NFIL3
MAFK
CEBPD
CEBPB
CEBPE
JDP2
CREB3
XBP1
DBP
CEBPG
ATF4
Cebpb
CREB3L1
Mafb
Dbp
1 2 11 3 4 5 6 7 8 9 10 12 13
1 2 3 4 5 6 7 8 9 11 12 13 10
1 2 11 3 4 5 6 7 8 9 10 12 13
DNA-Flexibility
0 1 2 3

Figure 5

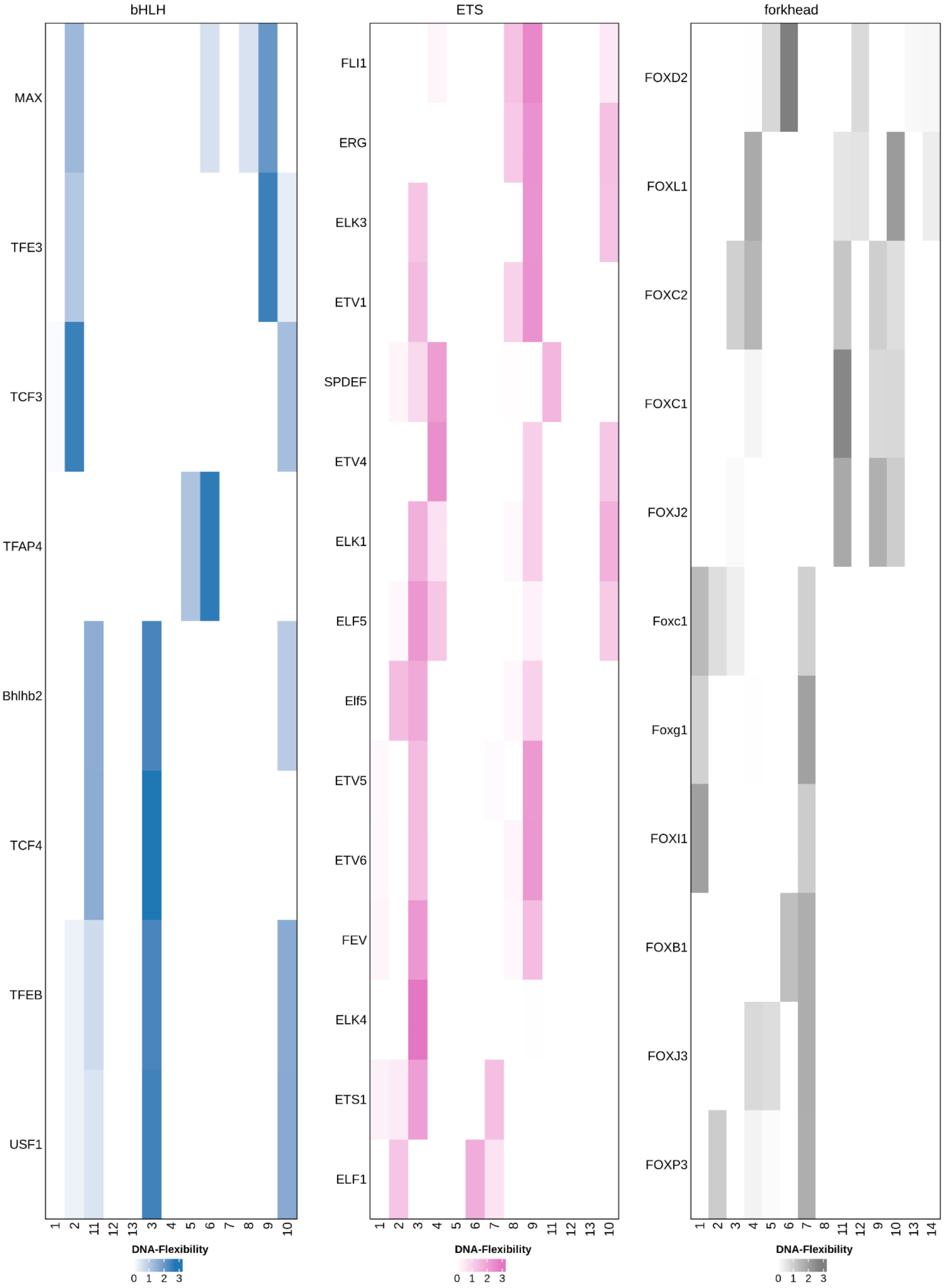
bHLH
MAX
TFE3
TCF3
TFAP4
Bhlhb2
TCF4
TFEB
USF1
1 2 11 12 13 3 4 5 6 7 8 9 10
DNA-Flexibility
0 1 2 3
ETS
FLI1
ERG
ELK3
ETV1
SPDEF
ETV4
ELK1
ELF5
Elf5
ETV5
ETV6
FEV
ELK4
ETS1
ELF1
1 2 3 4 5 6 7 8 9 11 12 13 10
DNA-Flexibility
0 1 2 3
forkhead
FOXD2
FOXL1
FOXC2
FOXC1
FOXJ2
Foxc1
Foxg1
FOXI1
FOXB1
FOXJ3
FOXP3
1 2 3 4 5 6 7 8 11 12 9 10 13 14
DNA-Flexibility
0 1 2 3

Figure 6

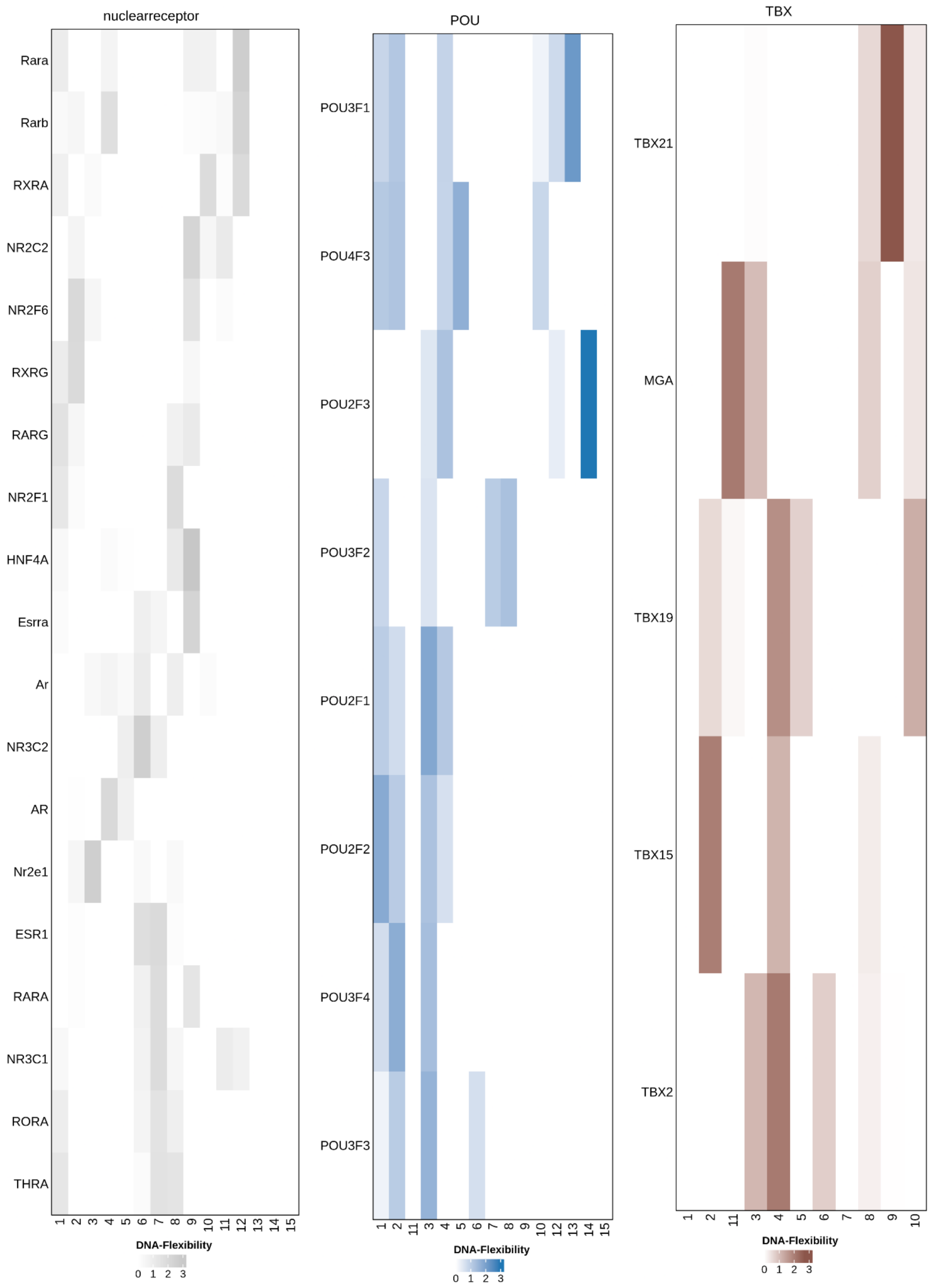
nuclearreceptor
Rara
Rarb
RXRA
NR2C2
NR2F6
RXRG
RARG
NR2F1
HNF4A
Esrra
Ar
NR3C2
AR
Nr2e1
ESR1
RARA
NR3C1
RORA
THRA
1 2 3 4 5 6 7 8 9 10 11 12 13 14 15
DNA-Flexibility
0 1 2 3
POU
POU3F1
POU4F3
POU2F3
POU3F2
POU2F1
POU2F2
POU3F4
POU3F3
1 2 11 3 4 5 6 7 8 9 10 12 13 14 15
DNA-Flexibility
0 1 2 3
TBX
TBX21
MGA
TBX19
TBX15
TBX2
1 2 11 3 4 5 6 7 8 9 10
DNA-Flexibility
0 1 2 3

Figure 7

**Supplementary Figure 3A-C. DNA Flexibility Profiles Across TF Families Reveal Family-Specific Indirect Readout Patterns.** Heatmaps display DNA flexibility contributions at individual nucleotide positions within TF binding sites for different transcription factor families: **(A)** Homeodomain (*left*), C2H2 zinc finger (*middle*), and bZIP (*right*); **(B)** bHLH, ETS, Forkhead; **(C)** nuclearreceptor, POU, TBX. Each row represents a distinct TF, and columns correspond to nucleotide positions along the binding site. The intensity of colors indicates the magnitude of contribution from DNA flexibility features to binding specificity, with darker shades denoting stronger influence. For the homeodomain family, hierarchical clustering grouped TFs into seven distinct clusters (dendogram not shown) reflecting diverse flexibility readout mechanisms within a TF-family. C2H2 and bZIP families show characteristic flexibility signatures at specific nucleotide positions, highlighting the family-specific nature of indirect readout mechanisms mediated by DNA conformational properties.

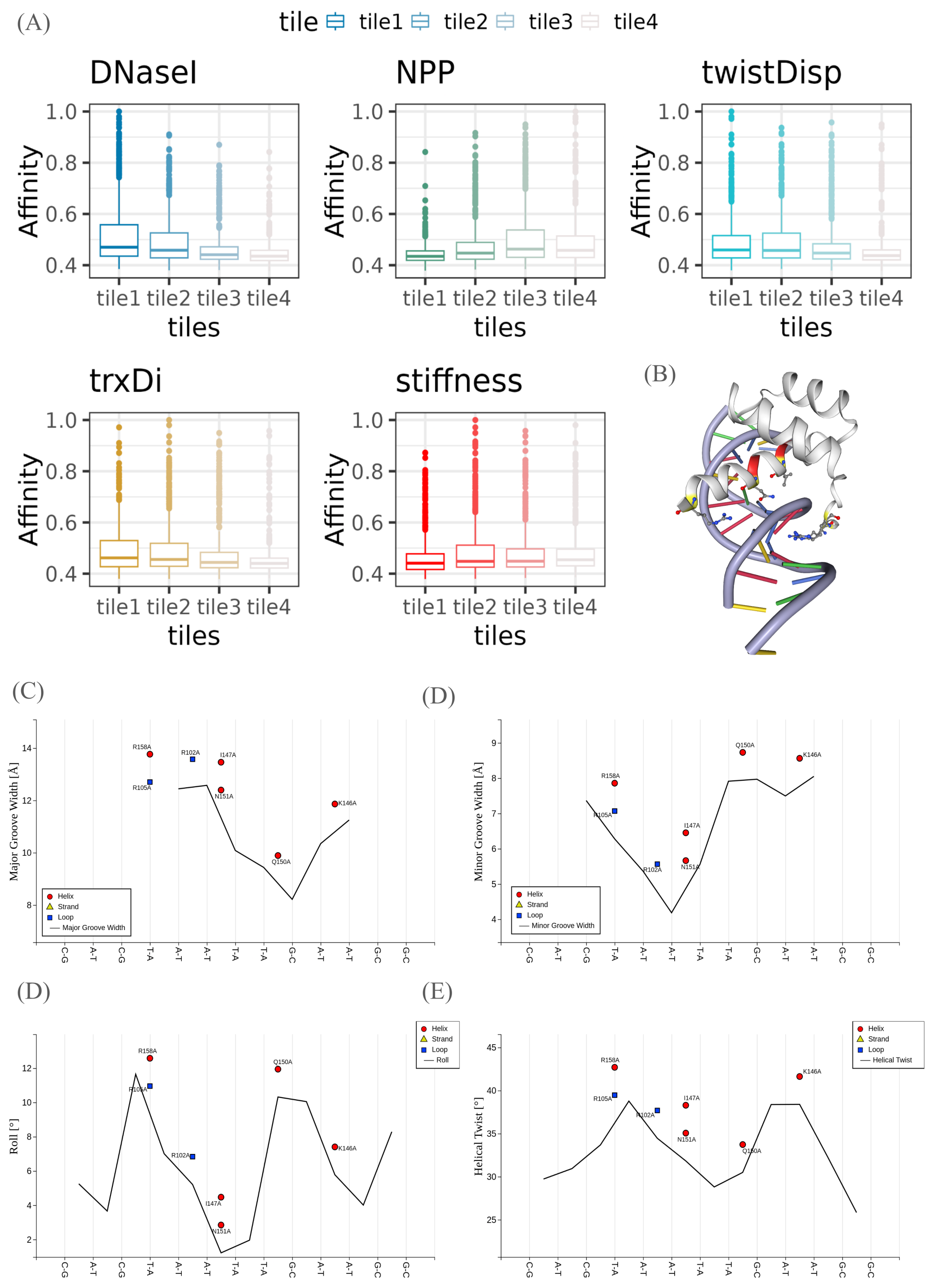
(A)
tile tile1 tile2 tile3 tile4
DNaseI
NPP
twistDisp
trxDi
stiffness
Affinity
tiles
tile1 tile2 tile3 tile4
(B)
(C)
Major Groove Width [Å]
Helix
Strand
Loop
Major Groove Width
(D)
Minor Groove Width [Å]
Minor Groove Width
(D)
Roll [°]
Roll
(E)
Helical Twist [°]
Helical Twist
R158A
R105A
R102A
I147A
N151A
Q150A
K146A
C-G A-T C-G T-A A-T A-T T-A T-A G-C A-T A-T G-C G-C

Figure 8

**Supplementary Figure 4: DNA Flexibility Contributions and Structural Analysis of Homeodomain TF MSX1 Binding Sites.** **(A–E)** Comparison of MSX1 binding affinity across bins of binding sites distributed by average DNA flexibility. Tile 1 to Tile 4 represents sequences with decreasing values of average flexibility features. **(F)** Diagram of major and minor groove contacts formed by MSX1 side chains, derived from the crystal structure. **(G–J)** Structural parameters calculated from the MSX1–DNA co-crystal structure using the x3DNA software in DNAproDB web server, **(G)** major groove width, **(H)** minor groove width, **(I)** roll, and **(J)** helical twist. Positions of critical amino acid residues involved in DNA recognition (e.g., Arg158, Arg105, Ile147, Asn151, Gln150, Lys146) are annotated with different shapes corresponding to secondary structural elements (helix, strand, loop). These plots illustrate how specific DNA shape features correlate with protein–DNA contacts and binding specificity.

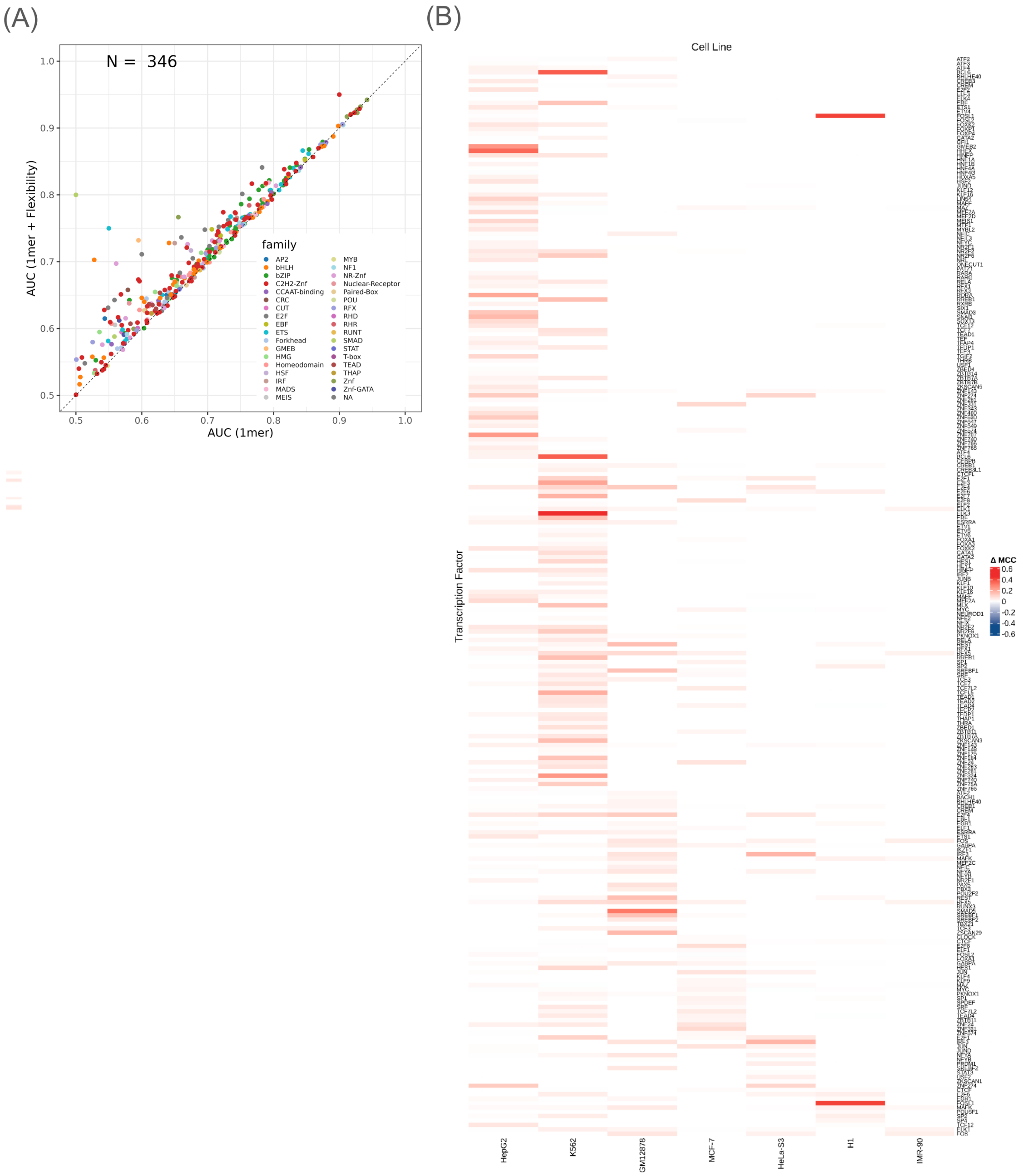
(A)
N = 346
AUC (1mer + Flexibility)
AUC (1mer)
0.5
0.6
0.7
0.8
0.9
1.0
family
AP2
bHLH
bZIP
C2H2-Znf
CCAAT-binding
CRC
CUT
E2F
EBF
ETS
Forkhead
GMEB
HMG
Homeodomain
HSF
IRF
MADS
MEIS
MYB
NF1
NR-Znf
Nuclear-Receptor
Paired-Box
POU
RFX
RHD
RHR
RUNT
SMAD
STAT
T-box
TEAD
THAP
Znf
Znf-GATA
NA
(B)
Cell Line
Transcription Factor
HepG2
K562
GM12878
MCF-7
HeLa-S3
H1
IMR-90
Δ MCC
0.6
0.4
0.2
0
-0.2
-0.4
-0.6

Figure 9

**Supplementary Figure 5. Cell Line-Specific Impact of DNA Flexibility on In Vivo TF Binding Prediction. (A)** Scatter plot comparing Area Under the Curve (AUC) for flexibility-augmented (1-mer + flexibility) versus sequence-only (1-mer) models across 355 high-confidence ENCODE ChIP-seq datasets. Each point represents a dataset; points above the diagonal indicate improved performance with flexibility features. **(B)** Heatmap illustrating the change in Matthews Correlation Coefficient ($\Delta$MCC) when DNA flexibility features are added to sequence-only models for predicting TF binding. Rows represent individual TFs, and columns represent different cell lines (K562, GM12878, HepG2, MCF-7, H1, HeLa-S3). Red shades indicate an improvement in MCC ($\Delta\text{MCC} > 0$) with the flexibility features. The intensity of the color corresponds to the magnitude of $\Delta$MCC, as shown in the legend.
